# Constrained non-coding sequence provides insights into regulatory elements and loss of gene expression in maize

**DOI:** 10.1101/2020.07.11.192575

**Authors:** Baoxing Song, Hai Wang, Yaoyao Wu, Evan Rees, Daniel J Gates, Merritt Burch, Peter J. Bradbury, Jeff Ross-Ibarra, Elizabeth A. Kellogg, Matthew B. Hufford, M. Cinta Romay, Edward S. Buckler

## Abstract

DNA sequencing technology has advanced so quickly, identifying key functional regions using evolutionary approaches is required to understand how those genomes work. This research develops a sensitive sequence alignment approach to identify functional constrained non-coding sequences in the Andropogoneae tribe. The grass tribe Andropogoneae contains several crop species descended from a common ancestor ~18 million years ago. Despite broadly similar phenotypes, they have tremendous genomic diversity with a broad range of ploidy levels and transposons. These features make Andropogoneae a powerful system for studying conserved non-coding sequence (CNS), here we used it to understand the function of CNS in maize. We find that 86% of CNS comprise known genomic elements e.g., *cis*-regulatory elements, chromosome interactions, introns, several transposable element superfamilies, and are linked to genomic regions related to DNA replication initiation, DNA methylation and histone modification. In maize, we show that CNSs regulate gene expression and variants in CNS are associated with phenotypic variance, and rare CNS absence contributes to loss of gene expression. Furthermore, we find the evolution of CNS is associated with the functional diversification of duplicated genes in the context of the maize subgenomes. Our results provide a quantitative understanding of constrained non-coding elements and identify functional non-coding variation in maize.

## Introduction

The genomes of a million eukaryote species could be sequenced within the next decade (Lewin et al. 2018). Yet understanding how those genomes work without ENCODE scale projects for each species will require that we use evolutionary approaches to identify key functional regions. DNA sequences subject to purifying selection across different species reflect functional constraints (Xiang et al. 2019; Finucane et al. 2015). The detection of conserved non-coding sequence (CNS) in plants is an ongoing challenge(Van de Velde et al. 2016) and recent studies(Tu et al. 2020; Ricci et al. 2019; Lu et al. 2019) could provide new insight into their function. In general, non-coding sequences occupy more of the genome than coding regions. The majority of GWAS hits are located in non-coding regions in maize and human etc, and enriched in gene expression regulators (Wallace et al. 2014; Nishizaki & Boyle 2017; Zhang & Lupski 2015; Giral et al. 2018), which have been extensively characterized recently (Lu et al. 2019; Ricci et al. 2019). Non-coding regions with high conservation(Haudry et al. 2013) are enriched with gene expression regulators (Polychronopoulos et al. 2017; Guo & Moose 2003; Vandepoele et al. 2006). However, links between CNS natural variation and expression variance have not been tested on a large scale in plants.

Genomes of the tribe Andropogoneae provide a valuable and powerful system for the study of constrained functional genomic elements. Andropogoneae species share a common ancestor around 16-20 million years ago(Vicentini et al. 2008), and, with their NADP-ME C4 (Sage & Zhu 2011; Black et al. 1969) photosynthesis, have become dominant warm-season grasses across the globe. Domesticated maize, sorghum, sugarcane, and miscanthus in the tribe are among the most productive crops in the world for grain, sugar, and biofuels (Brosse et al. 2012; Manners 2011), and many of the species look strikingly similar before flowering. This phenotypic stability belies tremendous genomic instability with multiple rounds of polyploidization and extremely active transposable elements (TEs) characterizing the tribe. Despite rapid sequence turnover elsewhere in Andropogoneae genomes, constrained non-coding sequences are potentially functional, making this an ideal clade for studying and clarifying their role.

## Results

### An atlas of conserved noncoding sequences in Andropogoneae tribe

Inspired by studies of regulatory architecture (Tu et al. 2020; Ricci et al. 2019), we use coding genes as anchors and develop a sensitive sequence alignment implementation to identify CNS in genomes with whole-genome duplications, numerous rearrangements, and gene loss(Schnable et al. 2011)(Fig. 1a). We aligned three released genomes (McCormick et al. 2018; Zhang et al. 2018; Swaminathan & Rokhsar) and two novel assembled genomes (Fig. 1b, Sfig. 1) against maize(Jiao et al. 2017). The sorghum genome is the only monoploid assembly without recent whole-genome duplication and generated the smallest CNS space (67.07 Mb, by counting alignment matches in maize genome). The largest CNS space was observed by aligning maize with sugarcane (86.97 Mb). The total number of CNS base-pairs present in maize and at least one other species was 106.52Mb, accounting for ~5% of the maize genome, hereafter, pan-And-CNS. A total of 42.27Mb sites were classified as conserved in all species and are called core-And-CNS. More species are needed to better describe pan- and core-And-CNS (Fig. 1c).

**Fig. 1.**
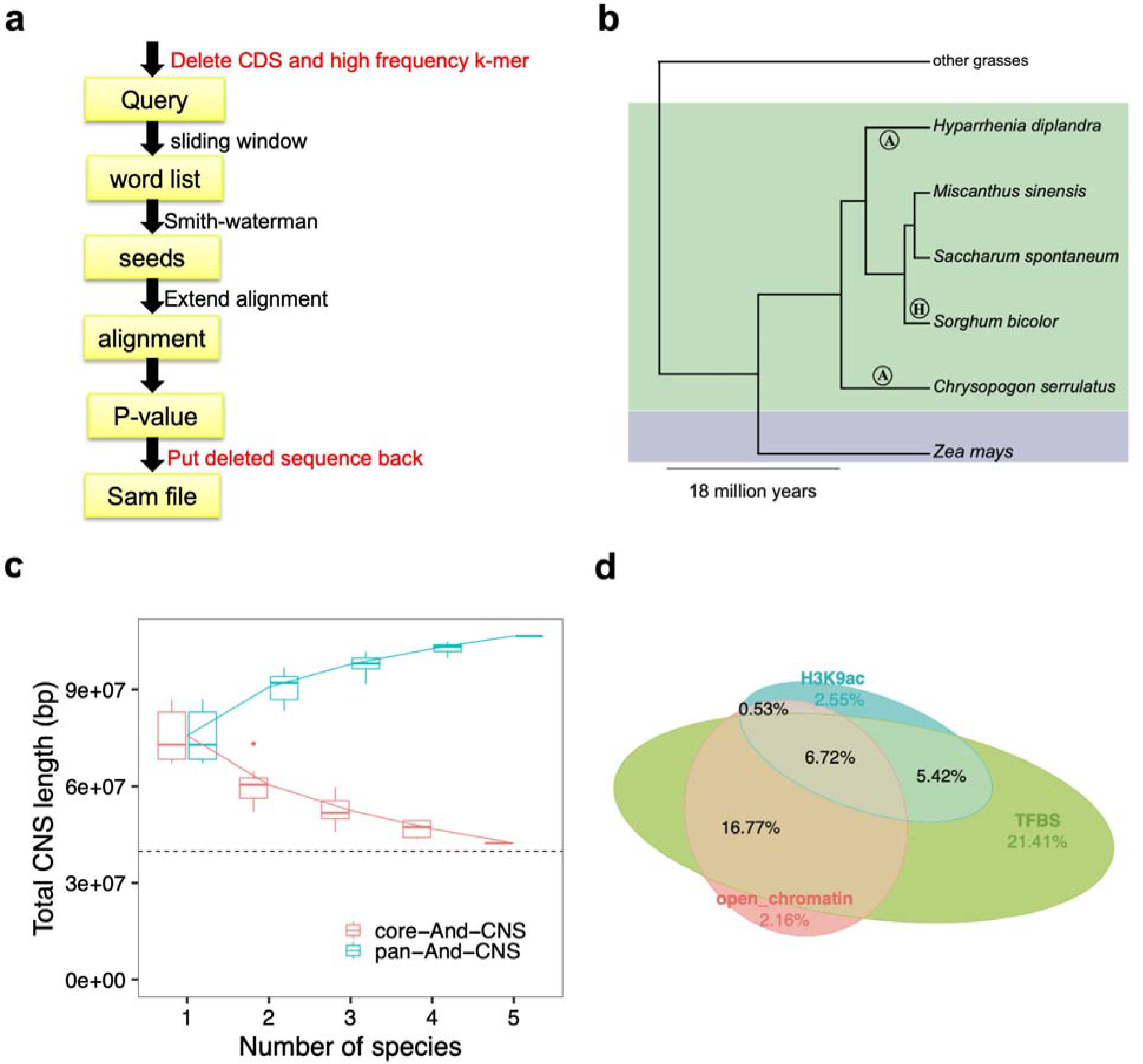
Only a partial Pan-Andropogoneae conserved non-coding sequence overlapped with *cis-*regulatory elements. **a,** The implemented algorithms to identify *Andropogoneae* conserved noncoding sequence using maize (*Zea mays L)* as reference. Briefly, we liftover the protein-coding gene annotation of maize APG V4 to other species by aligning the maize coding sequence (CDS) to other genomes using a splice site aware approach implemented in minimap2 [34]. The noncoding sequences of all other species were aligned against the masked maize genome using a k-mer match free, dynamic programming algorithm and optimized custom parameters. **b,** The phylogenetics topology of Andropogoneae species used in this study. The tree was constructed using plastome. The Sorghum *(Sorghum bicolor)* genome did not experience a recent wholegenome duplication (H). Maize and sorghum are two crop species. *Hyparrhenia diplandra,* and *Chrysopogon serrulatus* genomes were assembled in this project (A). The miscanthus (*Miscanthus sinensis*) genome is a diploid assembly and wild sugarcane (*Saccharum spontaneum L.)* is a tetraploid assembly. **c**, Simulations of the increase of the pan-And-CNS and the decrease of core-And-CNS base pairs by iteratively randomly sampling taxa. Solid lines indicate the pan- and core-And-CNS curves fit using points from all random combinations. The black horizontal dash line is the total base pairs of 10 maize chromosomes coding sequence (CDS). **d,** The proportion of intergenic pan-And-CNS overlapped with annotated regulatory elements.

### Not all conserved noncoding sequences overlapped with annotated *cis*-regulatory elements

We tested whether CNS are primarily known gene regulatory elements (Ricci et al. 2019; Tu et al. 2020). Since introns have a wide range of conserved functional roles (e.g., promote gene expression, guide splicing, non-coding RNA) (Chorev & Carmel 2012; Akua et al. 2010; Rigau et al. 2019; Greene et al. 1994; Ritchie & Flicek 2014), we did not sort these by function. Approximately half (52%) of the pan-And-CNS overlapped with coding genes (Sfig. 2), a percentage comparable to *Arabidopsis thaliana* (Haudry et al. 2013) and slightly higher than the proportion of genetic accessible chromatin in maize (Lu et al. 2019). Intergenic pan-And-CNS elements were 6 to 25 fold enriched for open chromatin regions (Ricci et al. 2019; Tu et al. 2020), transcription factor binding sites (TFBS)(Tu et al. 2020), and gene expression related Acetylation of histone 3 lysine 9 (H3K9ac) marks (Oka et al. 2017) (Sfig. 3), significantly higher than previous observation (Lai et al. 2017). Together these overlapped 25% of the pan-And-CNS (56% of intergenic pan-And-CNS) (Fig. 1d). A large proportion of genes have intergenic pan-And-CNS detected within 2Kb upstream, and they have higher expression levels and weaker tissue expression specification(Kadota et al. 2006) (Sfig. 4).

### Conserved non-coding sequence comprised a wide range of functional genomic elements

To uncover the functions of the rest of pan-And-CNS, we conducted a colocalization analysis of pan-And-CNS with annotated genomic elements (Fig. 2a). Chromosome contact domains were inferred to be conserved (Szabo et al. 2019). Hi-C technology characterized chromatin loops (Delaneau et al. 2019; Dong et al. 2017; Szabo et al. 2019), and their dynamic is associated with transcription activity(Dong et al. 2020). DNA interaction junctions identified by HI-C and HICHIP (Ricci et al. 2019; Peng et al. 2019) comprise a substantial extra proportion (21.23%) of intergenic pan-And-CNS. TE-derived CNSs have been reported by several independent studies (Dupeyron et al. 2019; Smith et al. 2008; Xie et al. 2006), although the contribution of TEs as regulatory elements is still controversial (de Souza et al. 2013; Zeng et al. 2018; Trizzino et al. 2018). We observed modest enrichment of CNS within 5 out of 13 TE superfamilies(Stitzer et al. 2019) (Fig. 2b), yet TEs only accounted for another 1.34% of the intergenic pan-And-CNS.

**Fig. 2.**
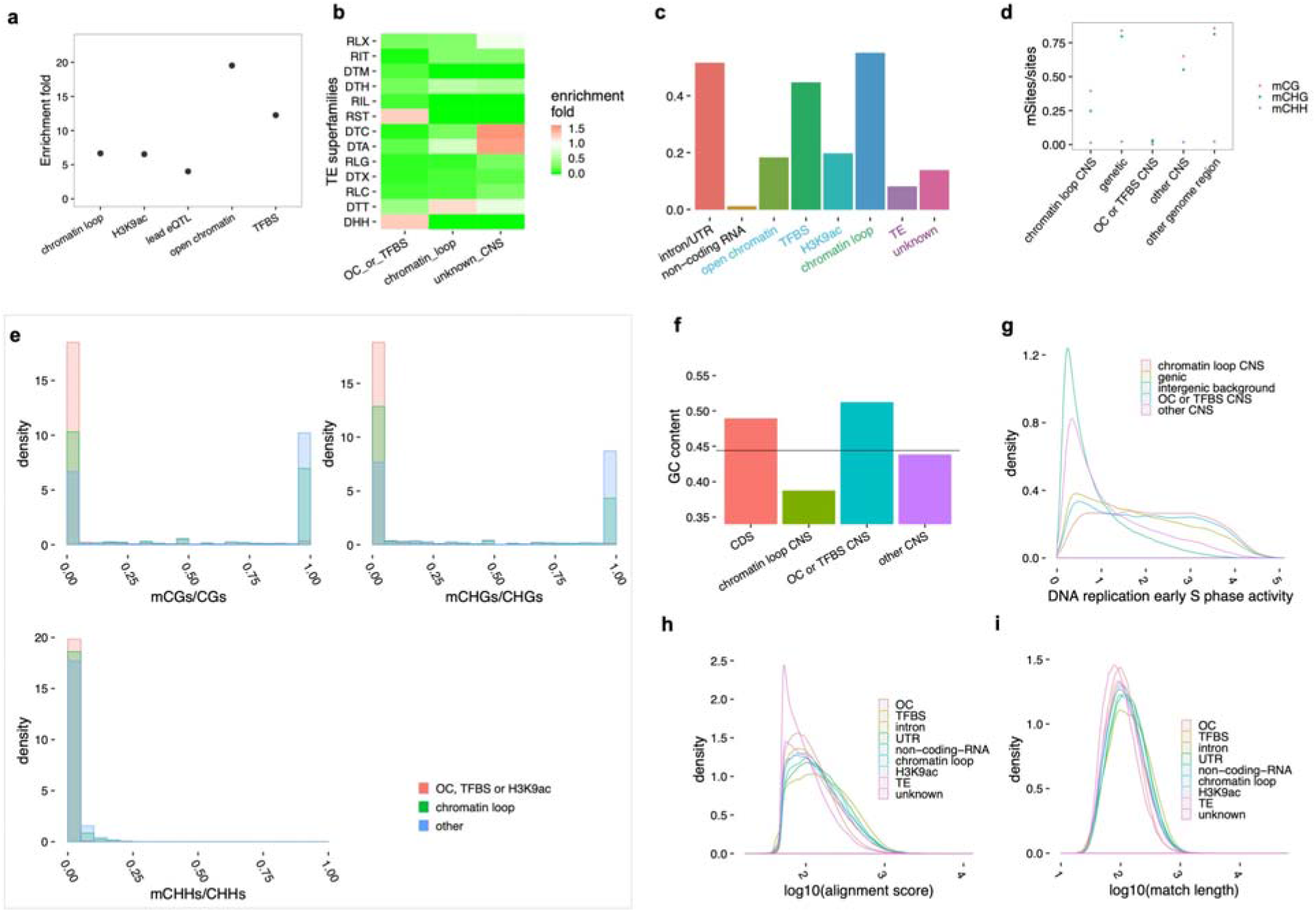
Pan-And-CNSs contribute to a wide range of nucleic molecular functions. **a,** Intergenic pan-And-CNSs are enriched in several types of *cis*-regulatory elements, lead eQTLs and chromatin structures. **b,** Enrichment was observed of different pan-And-CNSs groups in different TE superfamilies. (p value<0.0001, hypergeometric test) **c,** 86.28% of pan-And-CNS overlapped with diverse genomic elements. Only those 5 TE superfamilies with enrichment fold >1 were used here. Each CNS was allowed to overlap with multiple genomic elements. **d,e,** Different methylation ratios and patterns were observed for groups of pan-And-CNS. A subset of pan-And-CNSs did not overlap with known elements (other CNS) and have an intermediate methylation ratio(**d**). **f,** Groups of pan-And-CNSs have distinct GC content. The solid horizontal line is the average GC content of the masked maize genome. **g,** pan-And-CNSs are enriched in DNA replication initiation sites for the cell division cycle. **h,i,** The unexplained pan-And-CNSs have smaller sequence alignment scores (**h**) and shorter matched lengths (**i**). Within plots, OC is short for open chromatin.

We checked CNSs consisting of different function elements is due to incomprehensive genomic elements annotation or functionally diverse by investigating their sequence profiles. CNSs overlapping with open chromatin, TFBS or H3K9ac deposition sites (group A pan-And-CNS) have a low methylation ratio. This support the previous observation that *cis*-regulatory elements corresponding to low methylation ratio(Ricci et al. 2019). Whereas those in CNSs specifically overlapping with chromatin loops (group B) have a medium methylation ratio (Fig. 2d). GC and CHG methylation patterns (Ricci et al. 2019) further divide group B and unclassified pan-And-CNS into two distinguishable groups (Fig. 2e), this is consistent with previous reports that the A and B compartments of interphase chromosomes linked by Hi-C data are associated with distinct methylation(Dong et al. 2017). DNA methylation is associated with the evolution of GC content(Mugal et al. 2015). Group A pan-And-CNS have very high GC content, while group B pan-And-CNS have low GC content (Fig. 2f). DNA replication initiation regions(Wear et al. 2017) were associated with both group A and group B pan-And-CNSs (Fig. 2g). The observation of different sequences and methylation properties for CNS classes indicate that CNSs account for diverse functions.

Overall 14% of pan-And-CNS could not be attributed to any known function (Fig. 2c). The unknown CNSs are shorter and have lower sequence alignment scores (Fig. 2h,i). The overlapping enrichment fold with annotated elements of core-And-CNS is very similar to pan-And-CNS, with a slightly smaller proportion of unknown function (Fig. 2c, Sfig. 5). Considering the distribution of low methylation ratio and high DNA replication in unknown CNS regions (Fig. 2f,g), there may be some tissue-or species-specific(Lu et al. 2019) or uncharacterized functional elements that were not included in this study.

### Conserved non-coding sequence region comprise variants causing dysregulation

We further verified functional enrichment of CNS regions by testing if genotypic variants within the CNS region affect phenotype. Intergenic pan-And-CNS were enriched 4.02-fold (group A, 5.14; group B, 4.14) among the lead significant single nucleotide polymorphisms (SNPs) of expression genome-wide association study (eGWAS) in seven maize tissues(Kremling et al. 2018) (Fig. 2a). This is consistent with previous inferences that CNSs regulate gene expression(Meisler 2001). Using common CNS PAVs (with minor allele frequency>=0.1, 8.96%) as independent variables, we performed eGWAS using seven tissues (Fig. 3a). We found more than half of CNSs are located within 2.5Mb (2.5Mb is the estimated range of topologically associating domain (TAD) size (Gong et al. 2018), and almost all *cis*-regulatory elements are within 100Kb of nearest gene(Ricci et al. 2019; Tu et al. 2020)) of significantly associated genes (Fig. 3b, Sfig. 6, 7), this is consistent with enrichment overlapping between CNS with *cis-* regulatory element and interaction junctions. And CNS PAVs could be significantly associated with agronomic traits (Sfig. 8) directly.

**Fig. 3.**
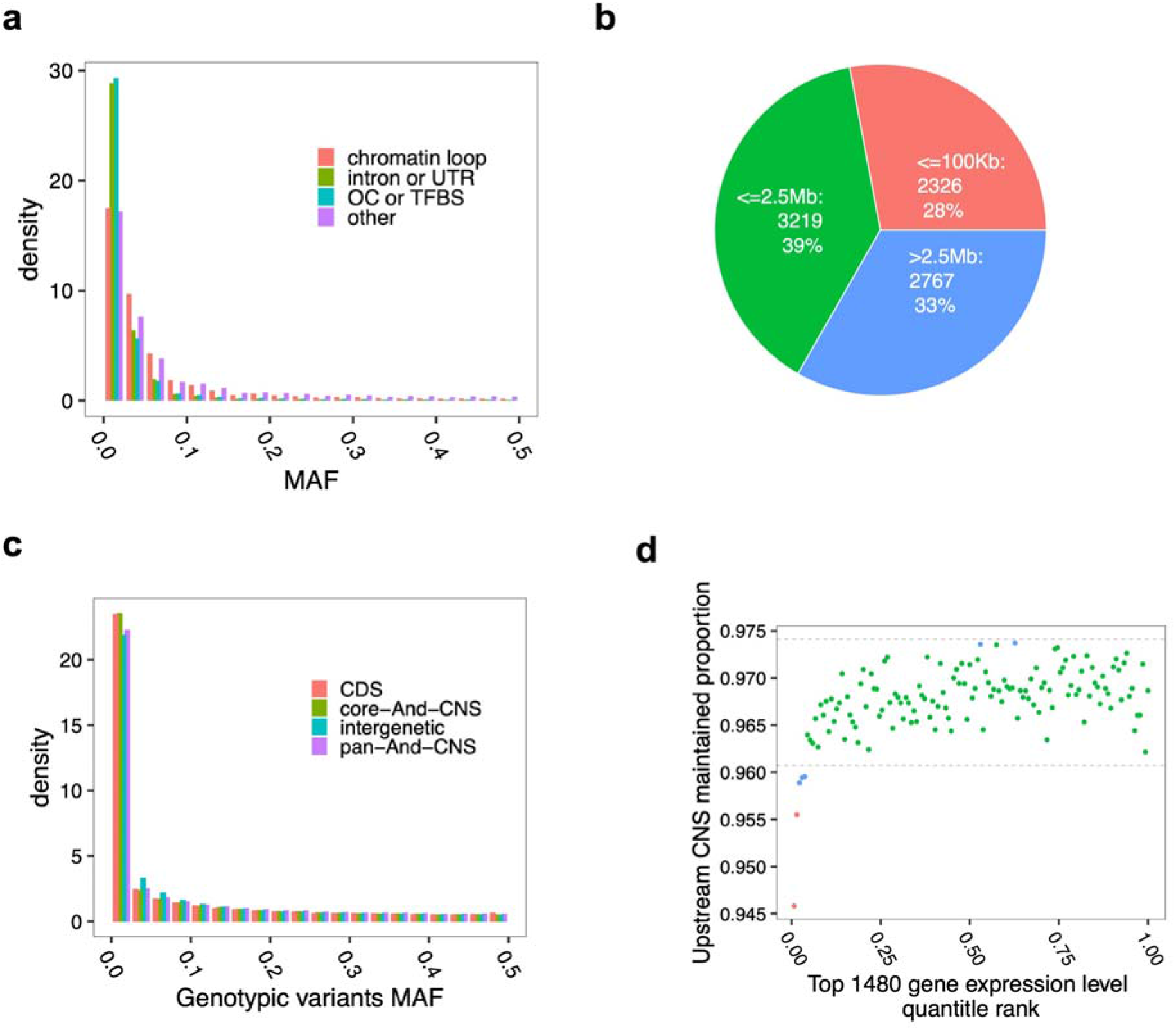
Variants in the CNS region contribute to gene expression variance. **a,** CNS PAVs can segregate at high MAF. A higher proportion of chromatin loop overlapping CNS and other non-*cis*-regulatory elements overlapping CNS with MAF > 0.1 were also observed. **b,** More than half of the significant associations of CNS PAVs with gene expression occur within 2.5Mb. The CNSs were encoded as binary variables and associated with gene expression level at the whole genome level. **c,** A higher proportion of rare variants was observed in CNS regions compared to the whole genome intergenic background. **d,** Comparing the proportion of maintained CNS in upstream 2Kb region of each accession and each gene with the gene expression level rank. Random shuffling the gene expression rank and CNS maintained proportion 1000 times, 99% (dash lines in gray) random intervals were calculated. Blue dots are out of 99% random intervals, and red dots are out of random intervals. CNS presence proportion is measured by counting the number of base-pairs. The gene expression level was quantified using maize agp v3 reference [53]. The top expressed 1500 genes of agp v3 were selected for this analysis and uplifted to the maize agp v4 reference [25]. This pattern is consistent across different tissues (Sfig. 10).

In a maize GWAS panel(Kremling et al. 2018), CNS region variants were not selected during recent domestication and local adaptation (Sfig. 9). And CNS regions(Kremling et al. 2018) have a larger proportion of variants with low minor allele frequency (MAF) and are enriched with rare variants (Fig. 3c). This is likely due to variants within CNS regions being under stronger purifying selection. Previous studies suggested rare variants contribute to dysregulation and are negatively associated with organism fitness (Flint-Garcia et al. 2005; Kremling et al. 2018). We further tested if CNS PAV is associated with extreme gene expression. Association between loss-of upstream (2Kb) CNS and extremely low expression rank of highly expressed genes was observed (Fig. 3d, Sfig. 10, 11).

### CNS variation is associated with functional diversity between Maize Subgenomes

CNS variation could also be observed among paralogous genes, which is common in maize because of a whole-genome duplication between 5 and 12 million years ago (Swigonová et al. 2004). We studied whether CNS evolution is associated with expression dosage of duplicated genes. We calculated the proportion of pan-And-CNS sites present in the upstream 2Kb of each syntenic duplicated pair (method) (Sfig. 12) and found a tendency that one CNS copy was maintained as complete while fragmented in the other copy (Fig. 4a and Sfig. 13), which is in contrast with the pattern of CDS (Sfig. 14). While the diversity of CNS is positively associated with CDS (Sfig. 15). Moreover, the orthologous copy with a larger proportion of pan-And-CNS maintained is associated with higher expression (Fig. 4b, Sfig. 16) (p-value=3e-15, Wilcoxon rank-sum test), which may be partially due to pseudogenization. The proportion of pan-And-CNS shared between syntenic paralogs is positively associated (r2=0.10, p-value< 2.2e-16) with expression similarity (measured using Pearson correlation coefficient) of duplicated genes (Sfig. 17, 18). These results suggest that CNS variation is associated with expression bias and functional diversity (subfunctionalization, neofunctionalization or pseudogenization) of duplicated genes.

**Fig. 4.**
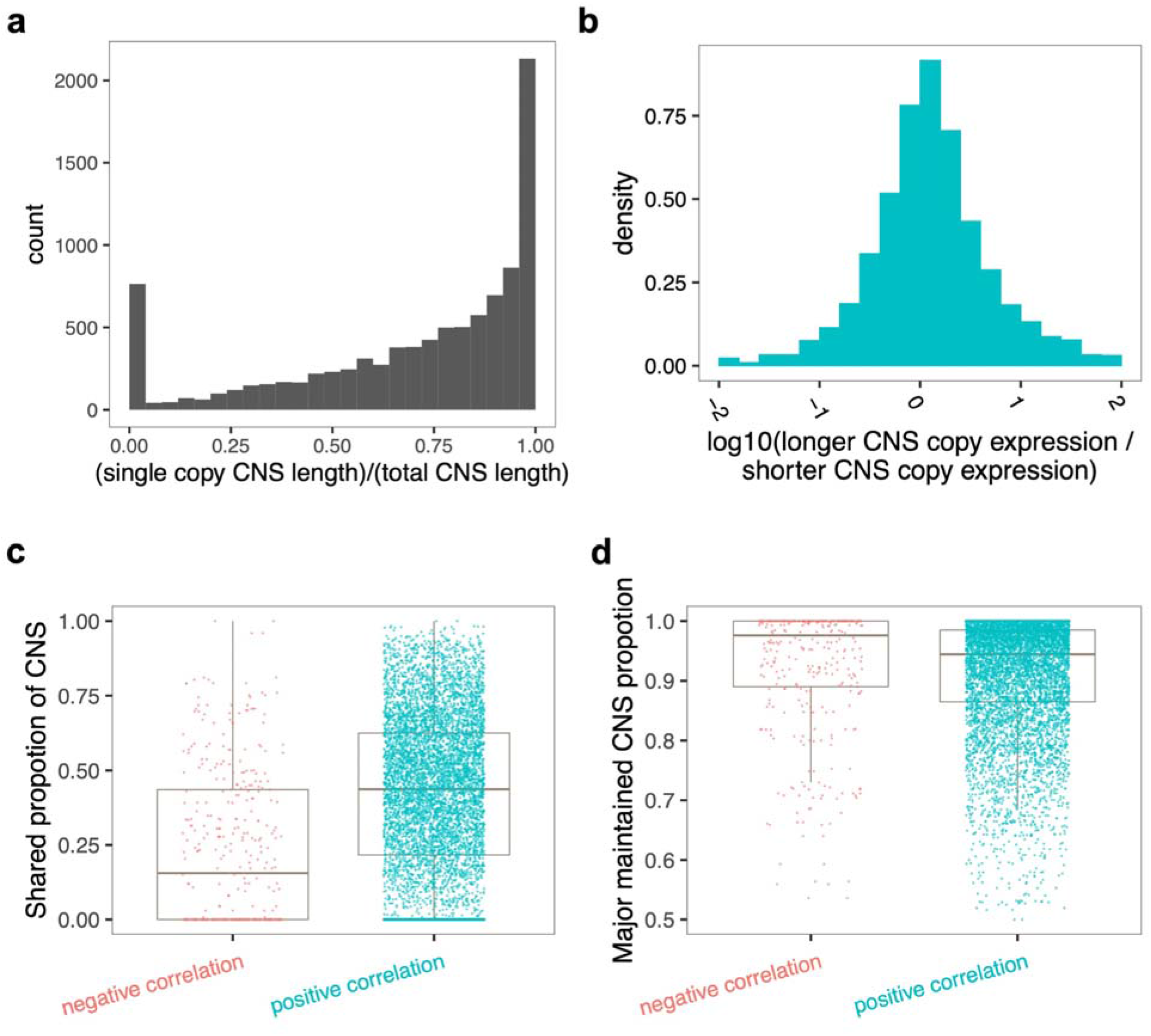
CNS diversity is associated with expression variance of paralogous genes. CNS detected using sorghum are showing here, and the duplicated genes pairs were detected by aligning maize with sorghum. **a,** In the upstream 2Kb region of homologous gene pairs from different subgenomes of maize, we counted how many base-pairs of CNS is present in each copy, how many base-pairs are shared and how many base-pairs in total. **b,** Between homologous gene pairs from different subgenomes of maize, the copy with longer CNS is associated with slightly higher expression level. Median values of expression levels across different tissues are shown here. **c,** Homologous gene pairs with negative expression correlation patterns share a smaller proportion of CNS compared to those pairs with positive expression correlation patterns. **d,** The major copy of homologous gene pairs with negative expression correlation maintained a larger proportion of CNS compared with the major copy of homologous gene pairs with positive expression correlation patterns. The major copy was defined as the one with longer CNS for this plot.

Duplicated pairs with negatively correlated expression levels shared a significantly smaller proportion of CNS than did positively correlated pairs (Fig. 4c) (p-value<2e-16, Wilcoxon rank-sum test). The copy with novel expression patterns may have new functions. The copy with longer CNS of the negatively correlated pairs typically had a higher proportion of the presumed ancestral set of CNSs compared to positively correlated pairs(Fig. 4d) (p-value=5e-10, Wilcoxon rank-sum test), indicating more similar with ancestral sequence.

## Discussion

Despite the challenges of working in genomes with vast numbers of transposons and duplications, a novel sensitive alignment approach shows that non-coding regions under purifying selection can be readily identified using evolutionary comparisons. This tool could identify CNS containing mismatches, indel and even TEs. Unlike previous CNS identification tools identifying very short CNS, our implementation considers the fact that sequence context of TFBS is important for its function(Tu et al. 2020). The CNSs identified here have a very high overlapping enrichment fold with TFBS etc.

The overlapping between CNS and *cis*-regulatory elements have been reported previously. Here we find CNS evolve a wide range of functions, i.e. chromatin loop, DNA replication. And their sequence properties are consistent with the functional classification. Those findings extended our knowledge of the function of non-coding sequences. By associating CNS PAV with gene expression level, we found a large proportion of distances between CNS PAV and distance significant genes are within the size of TAD, this reflects the impact of TAD structures on gene expression. By comparing gene expression and CNS diversity from two maize subgenomes, we also found loss-of CNS is mainly linked with decrease of gene expression. The evolution of CNS might be related with the dosage balance of gene expression, and associated with gene pseudogenization.

We found rare loss-of-CNS is associated with gene expression dysregulation, and mainly linked to loss-of-expression. Gene dysregulation has been associated with fitness decreasing. The CNS regions might provide targets for genome selection or genome editing for improving fitness of crops. As millions of species are sequenced, detailed alignment of thousands of species regulatory spaces will allow us to understand how each CNS is contributing to agronomic traits.

Overall 75% of the CNS are gene regulatory elements or introns, while many of the rest appear to either be involved in high order chromosome structure and perhaps DNA replication. Rare alleles have been shown to play an important role in expression(Kremling et al. 2018), but this study has shown the rare loss of expression is frequently associated with loss of CNSs. In species like maize, which have undergone ancient duplications, there is a tendency for only one of the two copies to retain the full complement of CNS and regulatory elements, which likely results in pseudogenization.

## Methods

### Plant material

*Hyparrhenia diplandra* was collected in Kenya by Rémy Pasquet *(Pasquet 1126)* and *Chrysopogon serrulatus* was obtained from the USDA Germplasm Repository Information Network (PI 219580; seed originally from Pakistan). Both plants were grown in the greenhouse at the Donald Danforth Plant Science Center. Vouchers of flowering specimens were deposited at the herbarium of the Missouri Botanical Garden; full specimen data are available through www.tropicos.org.

### DNA preparation and sequencing

DNA was extracted from young leaf tissue and we used the Nanopore MinION platform to conduct long read sequencing at Institute of Biotechnology, Cornell University. DNA with size 20-80Kb was selected following the Blue Pippin protocol, and the selected DNA was cleaned up using AMPure XP beads. DNA repair and End-Prep was performed by using NEB enzyme kits. After adapter ligation, we used the MinKNOW software to perform quality control for the MinION sequence library. Sequencing was performed following the manufacturer’s manual.

Illumina sequencing was performed by Novogene. A total of 1.0μg DNA per sample was used as input material for the DNA sample preparation. Sequencing libraries were generated using NEBNext^®^ DNA LibraryPrep Kit following the manufacturer’s recommendations and indices were added to each sample. Genomic DNA was randomly fragmented to a size of 350bp by shearing. Then DNA fragments were end polished, A-tailed, and ligated with the NEBNext adapter for Illumina sequencing, and further PCR enriched by P5 and indexed P7 oligos. PCR products were purified (AMPure XP system) and the resulting libraries were analyzed for size distribution by an Agilent 2100 Bioanalyzer and quantified using real-time PCR. The qualified libraries were fed into Illumina NovaSeq sequencers after pooling according to their effective concentration and expected data volume.

### Genome assembly

We used the NanoPlot(De Coster et al. 2018) and porechop package (Wick) package to check the quality of MinION reads and clean up. MinION clean reads were assembled using Flye v1.4.2(Kolmogorov et al. 2019) with default parameters. The MinION reads were mapped to the assembly using minimap2 (Li 2018) with “-x map-ont” parameter, and racon v1.3.1(Vaser et al. 2017) was used to polish the assembly with default parameters. This assembly polishing using MinION reads was repeated three times. The MEM module of BWA(Li & Durbin 2009) v 0.7.17 (parameter -k11 -r10) was used to map short reads to the MinION polished assembly. The “markdup” command implemented in samtools (Li et al. 2009) was used to remove duplicated short reads and Pilon(Walker et al. 2014) v 1.23 “--diploid --fix bases’’ was used for error correction (Sfig. 1A). Assembly correction using short reads was repeated three times.

### Genome assembly evaluation

We used a dataset of 5592 genes shared by four of the six Andropogoneae genomes (maize, sorghum, miscanthus, sugarcane) and an outgroup species *Setaria italica.* Their CDS sequences were mapped to the new assemblies, using minimap2 with parameters “-ax splice -a -uf -C 1 -k 12 -P -t 12 --cs”. For mapped genes, we defined the minimum extent of flanking to evaluate the assembly continuity (Sfig. 1C).

- n1=start_position
- n2=contig_length – end_position
- minimum extent of flanking = minimum (n1, n2)

BUSCO v 3.1.0 with the parameter “-m geno -sp maize -f -r -l” and database liliopsida_odb10 was used to evaluate the assembly quality.

### Syntenic gene detection

We used the quota-alignment pipeline to find syntenic genes (Tang et al. 2011) between maize and sorghum. The CDS sequence alignment was performed using blastn with parameters “-outfmt 6 -strand plus -task blastn -evalue 5 -word_size 7 -max_target_seqs 1000”. We modified blast_to_raw.py of quotaalignment package to avoid filtering of duplicated genes. The parameters of quota_align.py were set as “-merge --Dm=20 --min_size=3”. And the parameter of quota_align.py was set as “1:2”.

### CNS identification pipeline

We listover the reference gene structure annotation to the query genome using minimap2. CDS sequences and high-frequency k-mer regions were masked in both the reference genome and query genome. The upstream and down-stream 100Kb sequence and intron sequence of protein-coding genes were extracted to perform pairwise sequence alignment. In this analysis, upstream and downstream are used relative to the translation start and stop site, respectively. The range is selected based on the previous observation that almost all regulatory elements are located within 100Kb of the nearest gene (Rodgers-Melnick et al. 2016; Tu et al. 2020).

For each pair of sequence segments from different species, we take the maize sequence as reference and sequence from another species as the query. We used a Smith-Waterman algorithm(Smith & Waterman 1981) to generate alignment seeds. Each seed may be aligned as multiple copies in the reference genome. Each surviving seed is extended using a semi-global dynamic programming algorithm and extension is terminated using the X-drop approach (Zhang et al. 1998). The dynamic programming score was reported as sequence alignment score. Any extended sequence alignment that passes the p-value(Karlin & Altschul 1990) threshold (0.1 in this study, calculated in each alignment fragment independently without considering multiple tests) was categorized as CNS and output to a sam file, on which downstream analysis could be conducted taking advantage of well-developed applications (Li et al. 2009; Thorvaldsdóttir et al. 2013). We only used those CNS located on maize chr1-chr10 for all the analyses conducted in this project. The longest absolutely conserved CNS across all the 6 species was 113bp (Sfig. 19), which is shorter than that found in mammals(Bejerano et al. 2004).

### Enrichment analysis of CNS

The output sam files were reformatted into bam format using samtools, and the “depth” command of samtools was used to check how many unique base-pairs were classified as conserved. We counted how many unique CNS base pairs overlap with open chromatin.

Enrichment fold was calculated as:

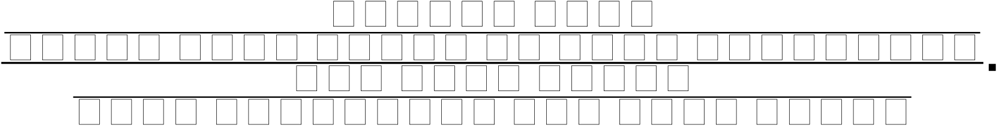

The enrichment overlapping with TFBS, H3K9ac enriched regions, eQTL, chromatin loop and TE was calculated in the same way.

### Overlap with maize non-coding RNA

lincRNA, miRNA or tRNA_gene annotated in the maize B73 v4 were used as non-coding RNA. And lncRNA annotated by Linqian et al(Han et al. 2019). was used to complement the B73 v4 annotation.

### Overlapping CNS with DNA duplication initial regions

The DNA duplication signals files were downloaded from the original publication(Wear et al. 2017). And the maize reference V3 coordinates were uplifted to V4 coordinates using CrossMap v0.2.8 (Zhao et al. 2014) and the chain file released from ensembl website. A density plot was created for the early phase signal of all the windows overlapped with each class of CNS or genetic element being investigated.

### CNS presence/absence variant encoding in maize population

The released Illumina short reads(Kremling et al. 2018) were mapped to maize APG v4 reference genome using “bwa mem” v 0.7.13 with default parameters. Each base pair was classified as “callable” and “noncallable” using GATK v 3.8(McKenna et al. 2010). The coverage for each base pair was checked using the “samtools mpileup’’ command. And reads were mapped to the CNS sequence using “bwa mem” with parameters “-a -c 200000 -S -P “. The coverage on the CNS sequence was also calculated using the”samtools mpileup” command. For each base pair of each CNS, if it is classified as “callable” or has coverage from genome reads mapping or CNS reads mapping, the base pair would be classified as “presence”, else “absence”. The CNS presence/absence was encoded by counting the CNS length and how many base pairs that were classified as “presence” for each CNS. And the above process was conducted for each resequenced maize accession independently.

Here only CNS detected using maize and sorghum were used, since it is easier to determine the start/end coordinate for each CNS fragment. And sorghum CNS, pan-And-CNS and core-And-CNS gave similar enrichment overlapping with open chromatin, TFBS and H3K9ac deposition sites. The absence of core-And-CNS are rare (Sfig. 20). Using all the pan-And-CNS, a lot of regions would be calculated multiple times. By linking all the overlapped pan-And-CNS, that would make it confusing about what is a CNS unit. Those taxa with low sequencing coverage were excluded (14Gbp of reads was used as a threshold), since they limited the CNS presence/absence variation (PAV) detection (Sfig. 21).

### CNS based association

For each CNS in each maize accession, we calculated a “present ratio” defined by “number of present base pair”/”CNS length”. If the present ratio >=0.8, it would be encoded as 1, if present ratio<=0.2, it would be encoded as 0, otherwise unknown.

To associate CNS PAV with gene expression, we used CNS PAVs with minor allele count >= 15 and known allele count >=35. The CNS PAVs were used as independent variables for the association. Accessions with both the target CNS and target gene deleted were excluded for the specific association test. Gene absence/presence variants were encoded in the same way as CNS. 25 PEER factors and 3 PCs(Kremling et al. 2018) calculated from a SNP-based kinship matrix were used as co-variants and association analysis was conducted using a fixed linear model. Association p-values were calculated using an f-test, with a significance threshold 1e-06. The association is performed using a homemade script which implements a linear association model and calculates p-value using f-test.

CNS PAVs were imputed in the NAM population using TASSEL 5; association was performed using the mixed model implemented, which dealt with missing genotypes by subsampling. 39310 CNS PAV with MAF >0.1 and minor allele count >20 were used to conduct the association analysis.

### Data access

All the CNS sam files are available in sam format at: https://figshare.com/s/6efc554fec80132fb9a6

The dCNS implementation is available at: https://github.com/baoxingsong/dCNS.

Further scripts for making plots are available at: https://github.com/baoxingsong/CNSpublication. The sequence reads and *de novo* assembly genome sequencing data is available at: ****.

## Acknowledgments

We thank Maria Katherine Mejia-Guerra for sharing the ATAC-seq and TBFS peak calling results and suggestions and comments on the open chromatin and TBFS analysis. Thank William F. Thompson for insights on the DNA replication phase. We thank Rémy Pasquet for providing plant material of *Hyparrhenia diplandra* and the USDA Germplasm Repository Information Network for material of *Chrysopogon serrulatus.* Thank Emre Cimen for the help of statistical analysis. Thank Guillaume Ramstein for talking about eQTL and GWAS analysis. Thank Robert Bukowski, Qi Sun and Arcadio Valdes Franco for the read mapping and variant calling of the maize population. Thank Sara Miller and Mingqiu Dai for proofreading the manuscript. The *Miscanthus sinensis* genome sequence was produced by the US Department of Energy Joint Genome Institute. This work used the Extreme Science and Engineering Discovery Environment (XSEDE), which is supported by National Science Foundation grant number ACI-1548562.

## Authors’ contributions

B.S. and E.S.B. designed the experiments and wrote the manuscript. B.S. implemented the dCNS software. B.S. and Y.W. identified CNS. B.S. and H.W. conducted maize subgenome analysis. B.S. and E.R. performed genome assembly. P.B. imputed CNS PAV in NAM population and performed GWAS together with B.S and M.B.. E.A.K. and M.C.R. prepared the sequencing materials. J.R.I. and D.J.G. conducted the domestication and local adaptation scan. J.R.I., E.A.K., and M.H. contributed ideas and provided technical help. All authors revised and reviewed the manuscript.

## References

Akua T, Berezin I, Shaul O. 2010. The leader intron of AtMHX can elicit, in the absence of splicing, low-level intron-mediated enhancement that depends on the internal intron sequence. BMC Plant Biol. 10:93.

Bejerano G et al. 2004. Ultraconserved elements in the human genome. Science. 304:1321–1325.

Black CC, Chen TM, Brown RH. 1969. Biochemical Basis for Plant Competition. Weed Sci. 17:338–344.

Brosse N, Dufour A, Meng X, Sun Q, Ragauskas A. 2012. Miscanthus: a fast-growing crop for biofuels and chemicals production. Biofuels Bioprod. Biorefin. 6:580–598.

Chorev M, Carmel L. 2012. The function of introns. Front. Genet. 3:55.

De Coster W, D’Hert S, Schultz DT, Cruts M, Van Broeckhoven C. 2018. NanoPack: visualizing and processing long-read sequencing data. Bioinformatics. 34:2666–2669.

Delaneau O et al. 2019. Chromatin three-dimensional interactions mediate genetic effects on gene expression. Science. 364. doi: 10.1126/science.aat8266.

Dong P et al. 2017. 3D Chromatin Architecture of Large Plant Genomes Determined by Local A/B Compartments. Mol. Plant. 10:1497–1509.

Dong P et al. 2020. Tissue-specific Hi-C analyses of rice, foxtail millet and maize suggest non-canonical function of plant chromatin domains. J. Integr. Plant Biol. 62:201–217.

Dupeyron M, Singh KS, Bass C, Hayward A. 2019. Evolution of Mutator transposable elements across eukaryotic diversity. Mob. DNA. 10:12.

Finucane HK et al. 2015. Partitioning heritability by functional annotation using genome-wide association summary statistics. Nat. Genet. 47:1228–1235.

Flint-Garcia SA et al. 2005. Maize association population: a high-resolution platform for quantitative trait locus dissection: High-resolution maize association population. Plant J. 44:1054–1064.

Giral H, Landmesser U, Kratzer A. 2018. Into the Wild: GWAS Exploration of Non-coding RNAs. Front Cardiovasc Med. 5:181.

Gong Y et al. 2018. Stratification of TAD boundaries reveals preferential insulation of superenhancers by strong boundaries. Nat. Commun. 9:542.

Greene B, Walko R, Hake S. 1994. Mutator insertions in an intron of the maize knotted1 gene result in dominant suppressible mutations. Genetics. 138:1275–1285.

Guo H, Moose SP. 2003. Conserved noncoding sequences among cultivated cereal genomes identify candidate regulatory sequence elements and patterns of promoter evolution. Plant Cell. 15:1143–1158.

Han L, Mu Z, Luo Z, Pan Q, Li L. 2019. New lncRNA annotation reveals extensive functional divergence of the transcriptome in maize. J. Integr. Plant Biol. 61:394–405.

Haudry A et al. 2013. An atlas of over 90,000 conserved noncoding sequences provides insight into crucifer regulatory regions. Nat. Genet. 45:891–898.

Jiao Y et al. 2017. Improved maize reference genome with single-molecule technologies. Nature. 546:524–527.

Kadota K, Ye J, Nakai Y, Terada T, Shimizu K. 2006. ROKU: a novel method for identification of tissue-specific genes. BMC Bioinformatics. 7:294.

Karlin S, Altschul SF. 1990. Methods for assessing the statistical significance of molecular sequence features by using general scoring schemes. Proc. Natl. Acad. Sci. U. S. A. 87:2264–2268.

Kolmogorov M, Yuan J, Lin Y, Pevzner PA. 2019. Assembly of long, error-prone reads using repeat graphs. Nat. Biotechnol. 37:540–546.

Kremling KAG et al. 2018. Dysregulation of expression correlates with rare-allele burden and fitness loss in maize. Nature. 555:520–523.

Lai X et al. 2017. STAG-CNS: An Order-Aware Conserved Noncoding Sequences Discovery Tool for Arbitrary Numbers of Species. Mol. Plant. 10:990–999.

Lewin HA et al. 2018. Earth BioGenome Project: Sequencing life for the future of life. Proc. Natl. Acad. Sci. U. S. A. 115:4325–4333.

Li H. 2018. Minimap2: pairwise alignment for nucleotide sequences. Bioinformatics. 34:3094–3100.

Li H et al. 2009. The Sequence Alignment/Map format and SAMtools. Bioinformatics. 25:2078–2079.

Li H, Durbin R. 2009. Fast and accurate short read alignment with Burrows-Wheeler transform. Bioinformatics. 25:1754–1760.

Lu Z et al. 2019. The prevalence, evolution and chromatin signatures of plant regulatory elements. Nat Plants. 5:1250–1259.

Manners JM. 2011. Chapter 3 -Functional Genomics of Sugarcane. In: Advances in Botanical Research. Kader, J-C & Delseny, M, editors. Vol. 60 Academic Press pp. 89–168.

McCormick RF et al. 2018. The Sorghum bicolor reference genome: improved assembly, gene annotations, a transcriptome atlas, and signatures of genome organization. Plant J. 93:338–354.

McKenna A et al. 2010. The Genome Analysis Toolkit: a MapReduce framework for analyzing next-generation DNA sequencing data. Genome Res. 20:1297–1303.

Meisler MH. 2001. Evolutionarily conserved noncoding DNA in the human genome: how much and what for? Genome Res. 11:1617–1618.

Mugal CF, Arndt PF, Holm L, Ellegren H. 2015. Evolutionary consequences of DNA methylation on the GC content in vertebrate genomes. G3. 5:441–447.

Nishizaki SS, Boyle AP. 2017. Mining the Unknown: Assigning Function to Noncoding Single Nucleotide Polymorphisms. Trends Genet. 33:34–45.

Oka R et al. 2017. Genome-wide mapping of transcriptional enhancer candidates using DNA and chromatin features in maize. Genome Biol. 18:137.

Peng Y et al. 2019. Chromatin interaction maps reveal genetic regulation for quantitative traits in maize. Nat. Commun. 10:2632.

Polychronopoulos D, King JWD, Nash AJ, Tan G, Lenhard B. 2017. Conserved non-coding elements: developmental gene regulation meets genome organization. Nucleic Acids Res. 45:12611–12624.

Ricci WA et al. 2019. Widespread long-range cis-regulatory elements in the maize genome. Nat Plants. 5:1237–1249.

Rigau M, Juan D, Valencia A, Rico D. 2019. Intronic CNVs and gene expression variation in human populations. PLoS Genet. 15:e1007902.

Ritchie GR, Flicek P. 2014. Computational approaches to interpreting genomic sequence variation. Genome Med. 6:87.

Rodgers-Melnick E, Vera DL, Bass HW, Buckler ES. 2016. Open chromatin reveals the functional maize genome. Proc. Natl. Acad. Sci. U. S. A. 113:E3177–84.

Sage RF, Zhu X-G. 2011. Exploiting the engine of C(4) photosynthesis. J. Exp. Bot. 62:2989–3000.

Schnable JC, Springer NM, Freeling M. 2011. Differentiation of the maize subgenomes by genome dominance and both ancient and ongoing gene loss. Proc. Natl. Acad. Sci. U. S. A. 108:4069–4074.

Schnable PS et al. 2009. The B73 maize genome: complexity, diversity, and dynamics. Science. 326:1112–1115.

Smith AM et al. 2008. A novel mode of enhancer evolution: the Tal1 stem cell enhancer recruited a MIR element to specifically boost its activity. Genome Res. 18:1422–1432.

Smith TF, Waterman MS. 1981. Identification of common molecular subsequences. J. Mol. Biol. 147:195–197.

de Souza FSJ, Franchini LF, Rubinstein M. 2013. Exaptation of transposable elements into novel cis-regulatory elements: is the evidence always strong? Mol. Biol. Evol. 30:1239–1251.

Stitzer MC, Anderson SN, Springer NM, Ross-Ibarra J. 2019. The Genomic Ecosystem of Transposable Elements in Maize. bioRxiv. 559922. doi: 10.1101/559922.

Swaminathan K, Rokhsar D. Miscanthus sinensis v7.1 DOE-JGI. Miscanthus sinensis v7.1 DOE-JGI. https://phytozome.jgi.doe.gov/pz/portal.html#!info?alias=Org_Msinensis_er.

Swigonová Z et al. 2004. Close split of sorghum and maize genome progenitors. Genome Res. 14:1916–1923.

Szabo Q, Bantignies F, Cavalli G. 2019. Principles of genome folding into topologically associating domains. Sci Adv. 5:eaaw1668.

Tang H et al. 2011. Screening synteny blocks in pairwise genome comparisons through integer programming. BMC Bioinformatics. 12:102.

Thorvaldsdóttir H, Robinson JT, Mesirov JP. 2013. Integrative Genomics Viewer (IGV): high-performance genomics data visualization and exploration. Brief. Bioinform. 14:178–192.

Trizzino M, Kapusta A, Brown CD. 2018. Transposable elements generate regulatory novelty in a tissue-specific fashion. BMC Genomics. 19:468.

Tu X et al. 2020. Decoding the regulatory architecture of the maize leaf. bioRxiv. 2020.01.07.898056. doi: 10.1101/2020.01.07.898056.

Vandepoele K, Casneuf T, Van de Peer Y. 2006. Identification of novel regulatory modules in dicotyledonous plants using expression data and comparative genomics. Genome Biol. 7:R103.

Van de Velde J, Van Bel M, Vaneechoutte D, Vandepoele K. 2016. A Collection of Conserved Noncoding Sequences to Study Gene Regulation in Flowering Plants. Plant Physiol. 171:2586–2598.

Vaser R, Sović I, Nagarajan N, Šikić M. 2017. Fast and accurate de novo genome assembly from long uncorrected reads. Genome Res. 27:737–746.

Vicentini A, Barber JC, Aliscioni SS, Giussani LM, Kellogg EA. 2008. The age of the grasses and clusters of origins of C 4 photosynthesis. Glob. Chang. Biol. 14:2963–2977.

Walker BJ et al. 2014. Pilon: an integrated tool for comprehensive microbial variant detection and genome assembly improvement. PLoS One. 9:e112963.

Wallace JG et al. 2014. Association Mapping across Numerous Traits Reveals Patterns of Functional Variation in Maize. PLoS Genetics. 10:e1004845. doi: 10.1371/journal.pgen.1004845.

Wear EE et al. 2017. Genomic Analysis of the DNA Replication Timing Program during Mitotic S Phase in Maize (Zea mays) Root Tips. Plant Cell. 29:2126–2149.

Wick R. Porechop. Github https://github.com/rrwick/Porechop (Accessed December 24, 2019).

Xiang R et al. 2019. Quantifying the contribution of sequence variants with regulatory and evolutionary significance to 34 bovine complex traits. Proc. Natl. Acad. Sci. U. S. A. 116:19398–19408.

Xie X, Kamal M, Lander ES. 2006. A family of conserved noncoding elements derived from an ancient transposable element. Proc. Natl. Acad. Sci. U. S. A. 103:11659–11664.

Zeng L, Pederson SM, Kortschak RD, Adelson DL. 2018. Transposable elements and gene expression during the evolution of amniotes. Mob. DNA. 9:17.

Zhang F, Lupski JR. 2015. Non-coding genetic variants in human disease. Hum. Mol. Genet. 24:R102–10.

Zhang J et al. 2018. Allele-defined genome of the autopolyploid sugarcane Saccharum spontaneum L. Nat. Genet. 50:1565–1573.

Zhang Z, Berman P, Miller W. 1998. Alignments without low-scoring regions. J. Comput. Biol. 5:197–210.

Zhao H et al. 2014. CrossMap: a versatile tool for coordinate conversion between genome assemblies. Bioinformatics. 30:1006–1007.

