## Supplemental for "Constrained non-coding sequence provides insights into regulatory elements and loss of gene expression in maize"

**Sfigures:**

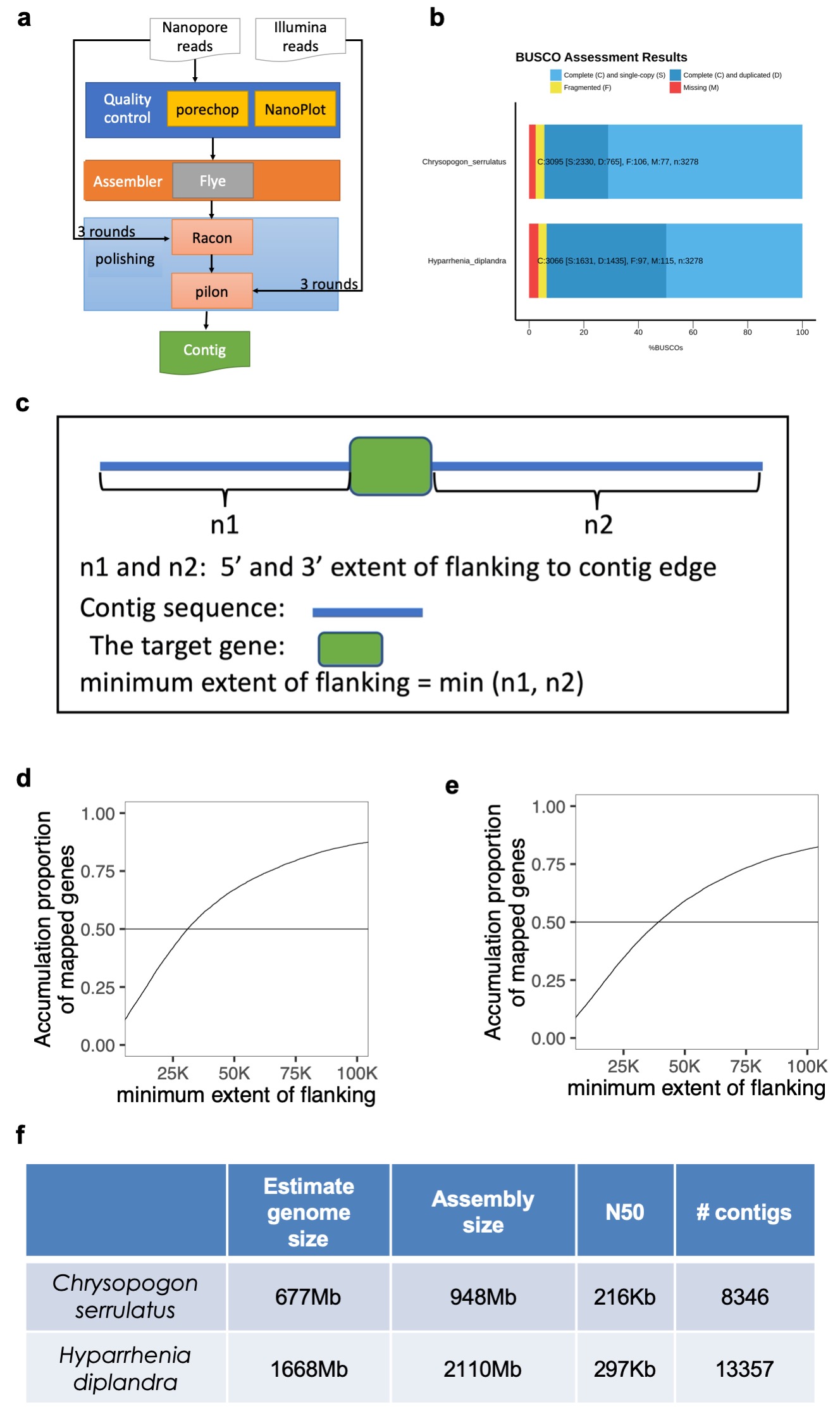

Sfig. 1 The genome assembly of *Chrysopogon serrulatus* and *Hyparrhenia diplandra*. **a,** The pipeline was used for genome assembly. **b,** The genome assembly complementness assessment using BUSCO pipeline[[72]](https://paperpile.com/c/mX0AVj/n7mX). **c,** A carton of minimum extent of flanking. **d, e,** The assembly continuity evaluated using the minimum extent of flanking for *Chrysopogon serrulatus*(d) and *Hyparrhenia diplandra*(e). **f,** The estimated genome size using flow cytometry approach and summary of assembly.

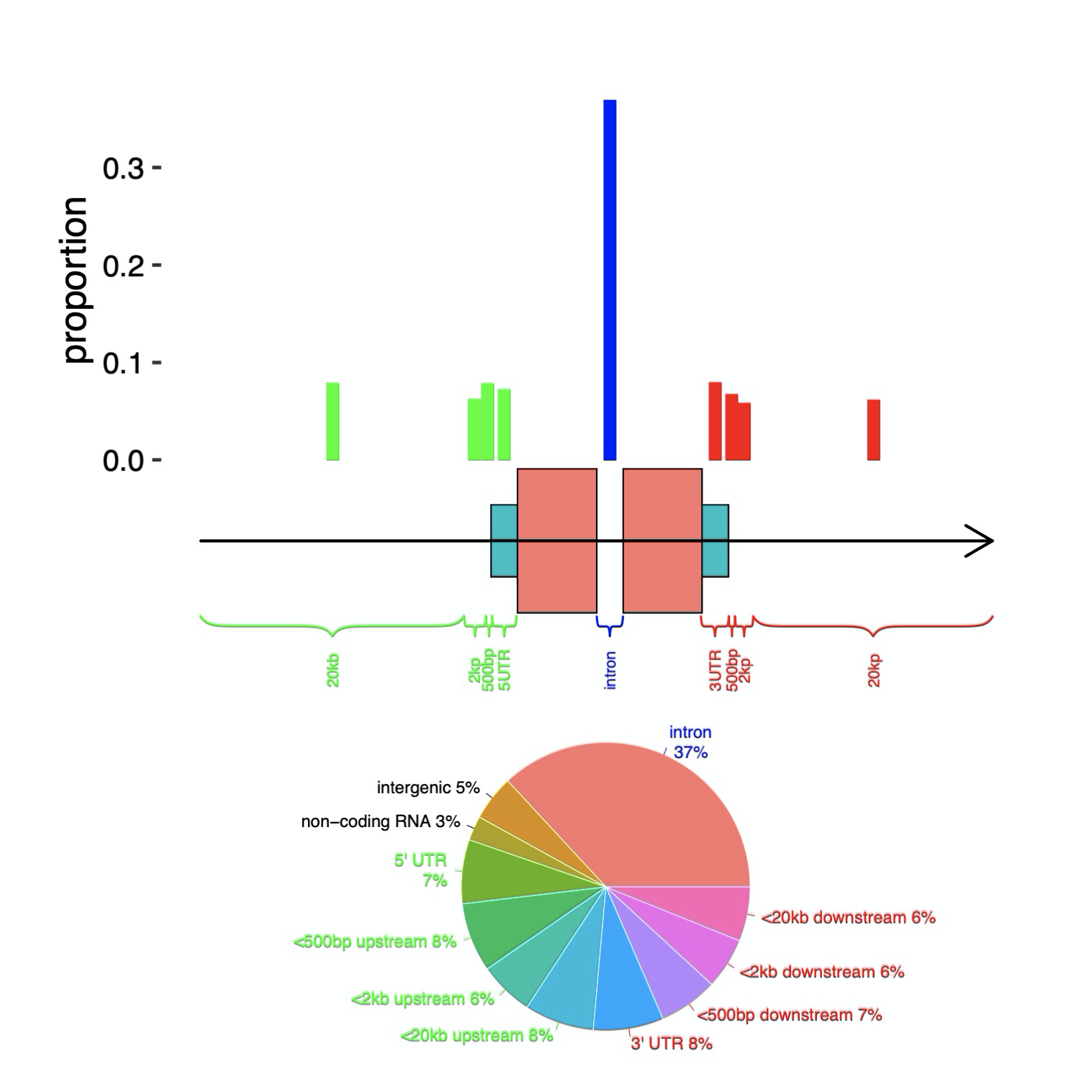

Sfig. 2 The distribution of detected CNS in different regions. 52% of the pan-And-CNS overlapped with intron or UTR of coding genes. 27% of pan-And-CNS were within 2Kb of genes (defined as overlapping ≥1 bp with the 2Kb regions flanking genes, but not overlapping the genes themselves). We also found 14% of pan-And-CNS occurred >2Kb from their nearest genes and 5% distal CNSs exceeded 20Kb from their nearest genes.

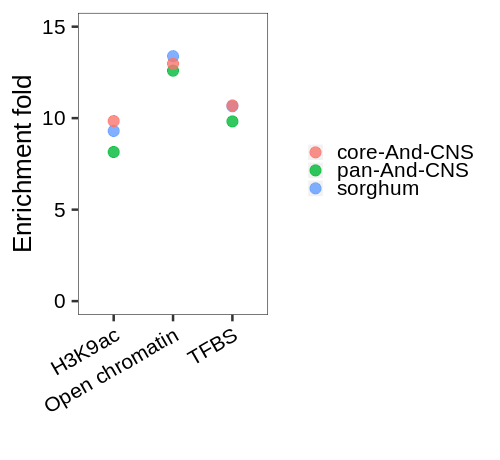

Sfig. 3 Pan-And-CNS, core-And-CNS and CNS detected by aligning maize with sorghum gave similar enrichment folds by overlapping with transcription factor binding sites, open chromatin region and H3K9ac deposition. This might be due to the equally phylogenetic distance between maize and another 5 species.

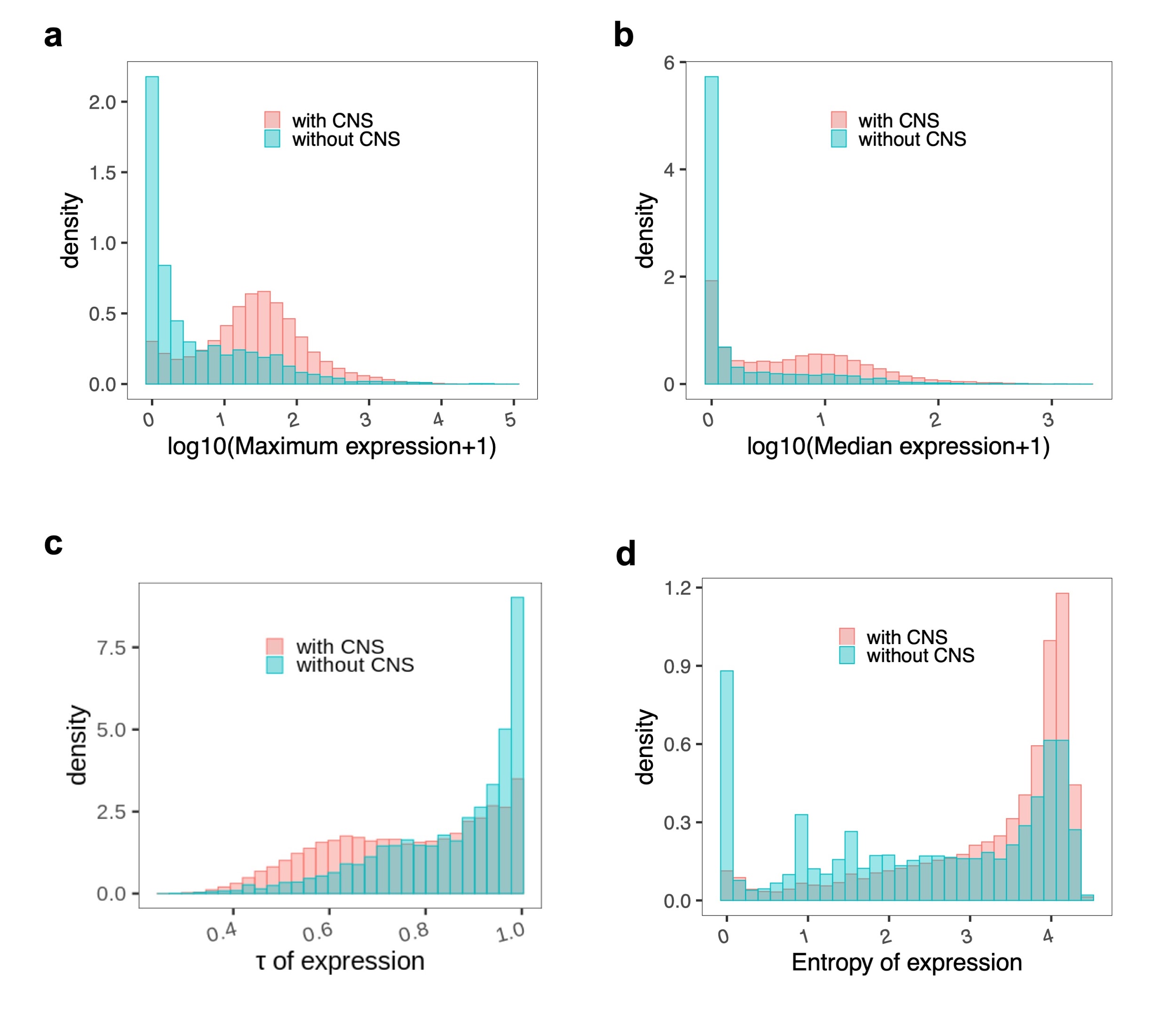

Sfig. 4 Those genes with pan-And-CNS detected within 2Kb of upstream have higher expression level and stronger tissues expression specification. Expression from 23 maize tissues[[73]](https://paperpile.com/c/mX0AVj/WWqM) were used here. There were 28950 genes that could be uniquely mapped between maize AGP v3 and AGP v4 annotation. 25127 of them have upstream CNS detected within 2Kb range. (a) Comparing the maximum gene expression across 23 tissues of genes with and without CNS detected within 2kb upstream. (b) Comparing median gene expression level. (c) Comparing tissue expression specification measured using τ[[74]](https://paperpile.com/c/mX0AVj/Gzj9). τ is valued between 0 for housekeeping genes and 1 for tissue-specific genes. (d) comparing tissue expression specification measured using entropy[[33]](https://paperpile.com/c/mX0AVj/diOZ), low entropy value is an indication of tissue expression specification.

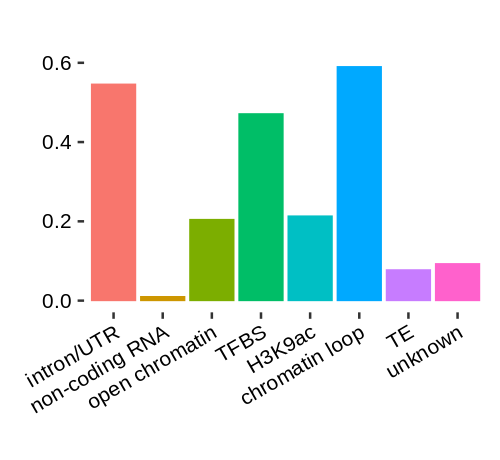

Sfig. 5 Proportion of core-And-CNS overlapped with defined elements. The core-And-CNSs were defined as CNSs present in all of the used species and have a length of at least 30 bp. We did not check sequence masking to define core-And-CNS. And a single core-And-CNS could overlap with several genomic elements. So the sum of all the proportions could be larger than 1.

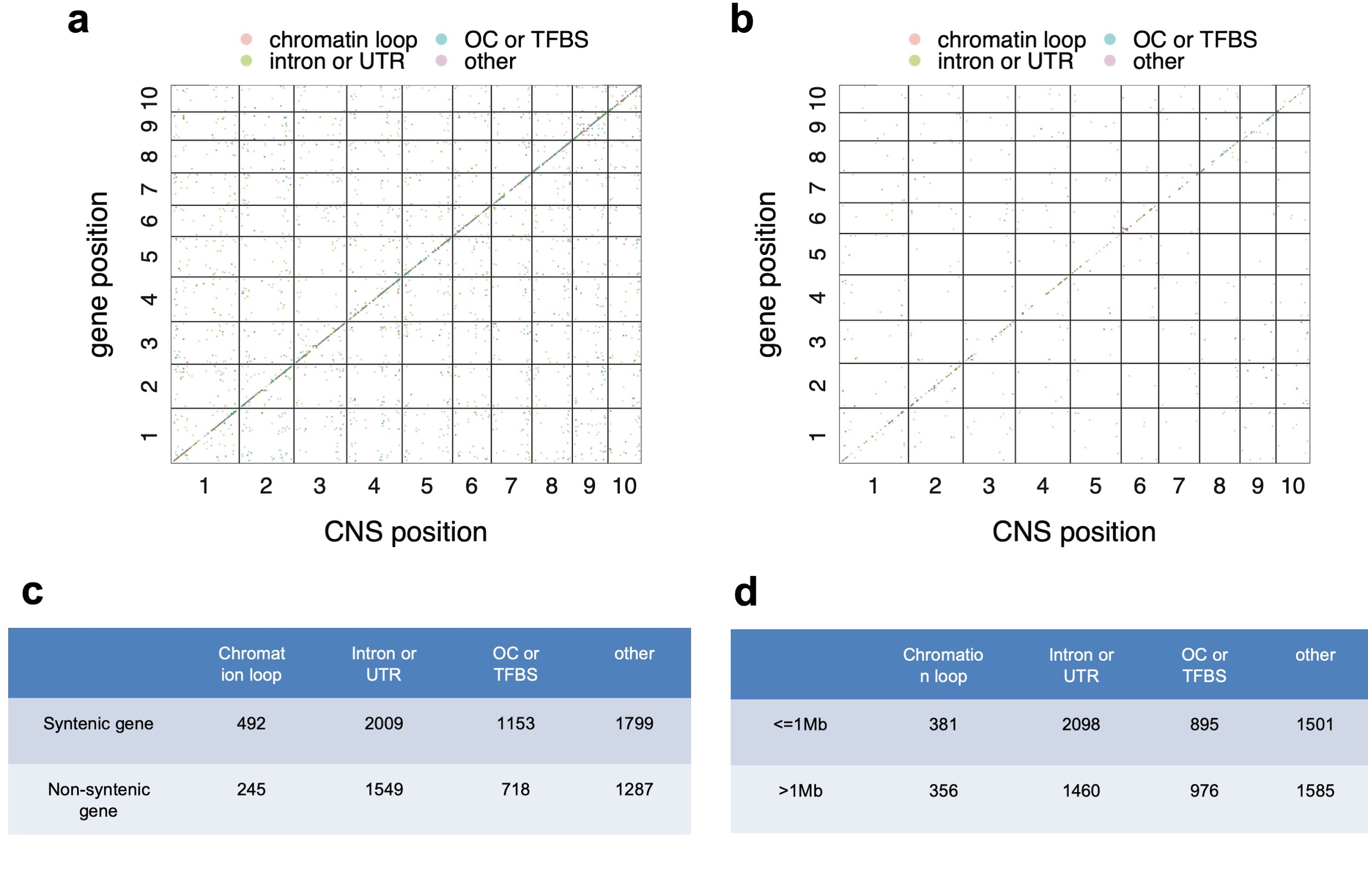

Sfig. 6 Summary of CNS PAV based maize eQTL analysis using root tissue classified by syntenic gene and non-syntenic gene and CNS classification. **a,** eQTL plot of syntenic genes. **b,** eQTL plot of non-syntenic genes.

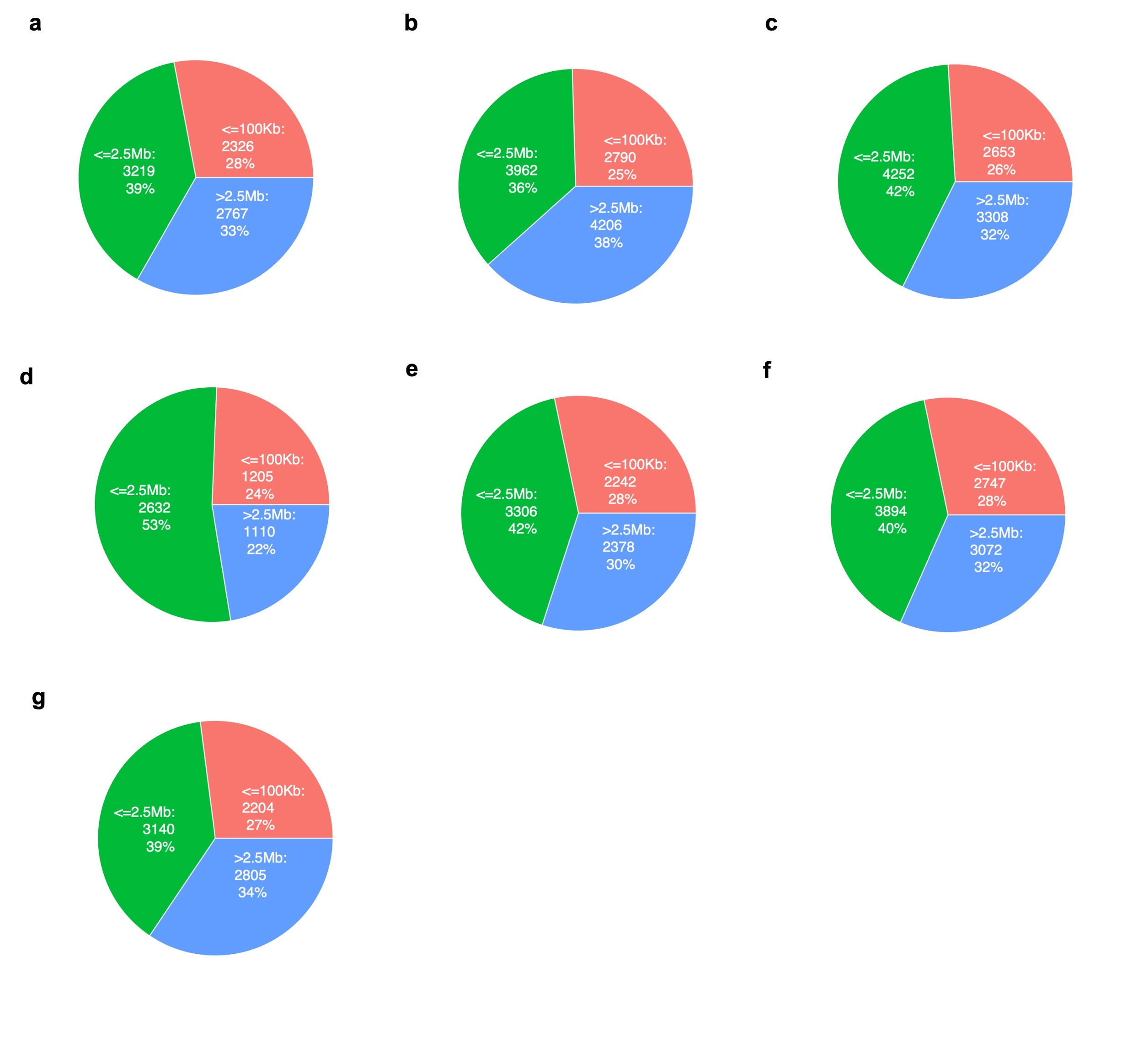

Sfig. 7 CNS PAVs regulate gene expression mainly in a cis- manner. **a,** Roots of germinating seedlings. **b**, Shoots of germinating seedling. **c**, Adult leaves collected at night. **d**, Adult leaves collected during the day. **e**, Tip of leaf three. **f**, Base of leaf three. **g**, Kernels at 350-growing-degree days.

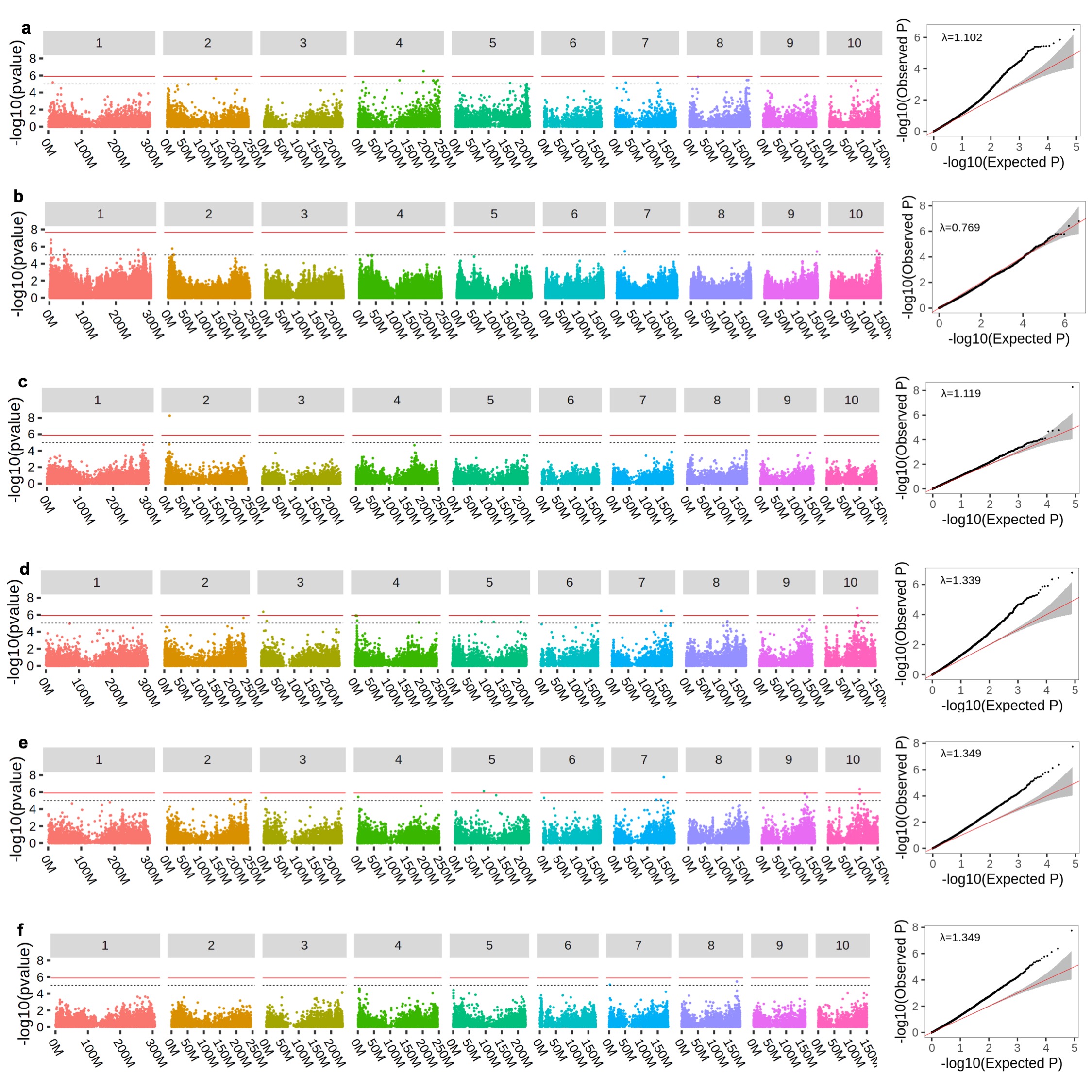

Sfig. 8 Manhattan and QQ plots of CNS PAV GWAS using the maize NAM population. The CNS PAV was called using NAM founder lines, and a custom TASSEL 5 plugin was implemented to impute CNS in the NAM population. The mixed linear model does not require genotype imputation for missing or unknown CNS PAVS, therefore this model could handle the ambiguity of the functional state of CNSs with intermediated amounts of missingness. **a**, chlorophyll a[[8]](https://paperpile.com/c/mX0AVj/dbxx). **b**, cob diameter[[75]](https://paperpile.com/c/mX0AVj/Xs45), the lead variant is located at Chr2:12902394, approximal to the candidate gene *zfl2*. **c,** cob diameter associated with SNPs. By keeping non-missing individuals of the lead CNS and lead SNPs on chromosome 2, the lead SNP and lead CNS have a LD with r2 value of 0.163. The lead SNP is located at chr2:11405490 and the lead CNS is located at chr2:12902394-12902454. The p-value of lead SNP is 3.67e-7 and the p-value of lead CNS is 5.08e-9. **d**, days to anthesis[[76]](https://paperpile.com/c/mX0AVj/mQXl). **e**, days to silk[[76]](https://paperpile.com/c/mX0AVj/mQXl). **f**, fumarate[[8]](https://paperpile.com/c/mX0AVj/dbxx).

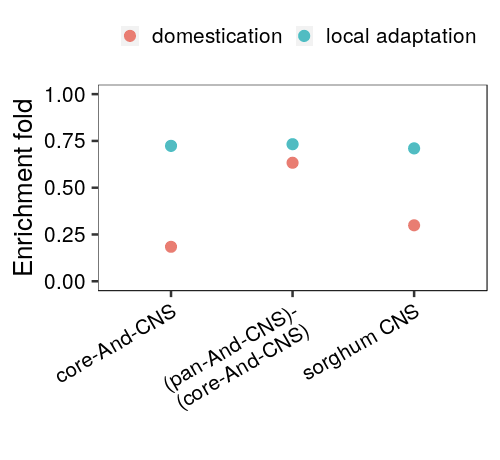

Sfig. 9 CNS regions are negatively enriched with domestication and local adaptation selected SNPs.

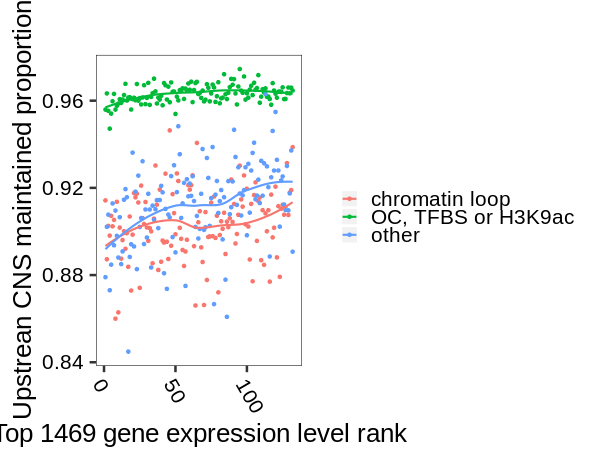

Sfig. 10 Loss-of-CNS is associated with a decrease in gene expression level. 1500 top expressed gene was selected using maize v3 reference and 1469 of them were be lifted to v4 annotation. The regression was plotted using the local regression function implemented in the R ggplot2 package.

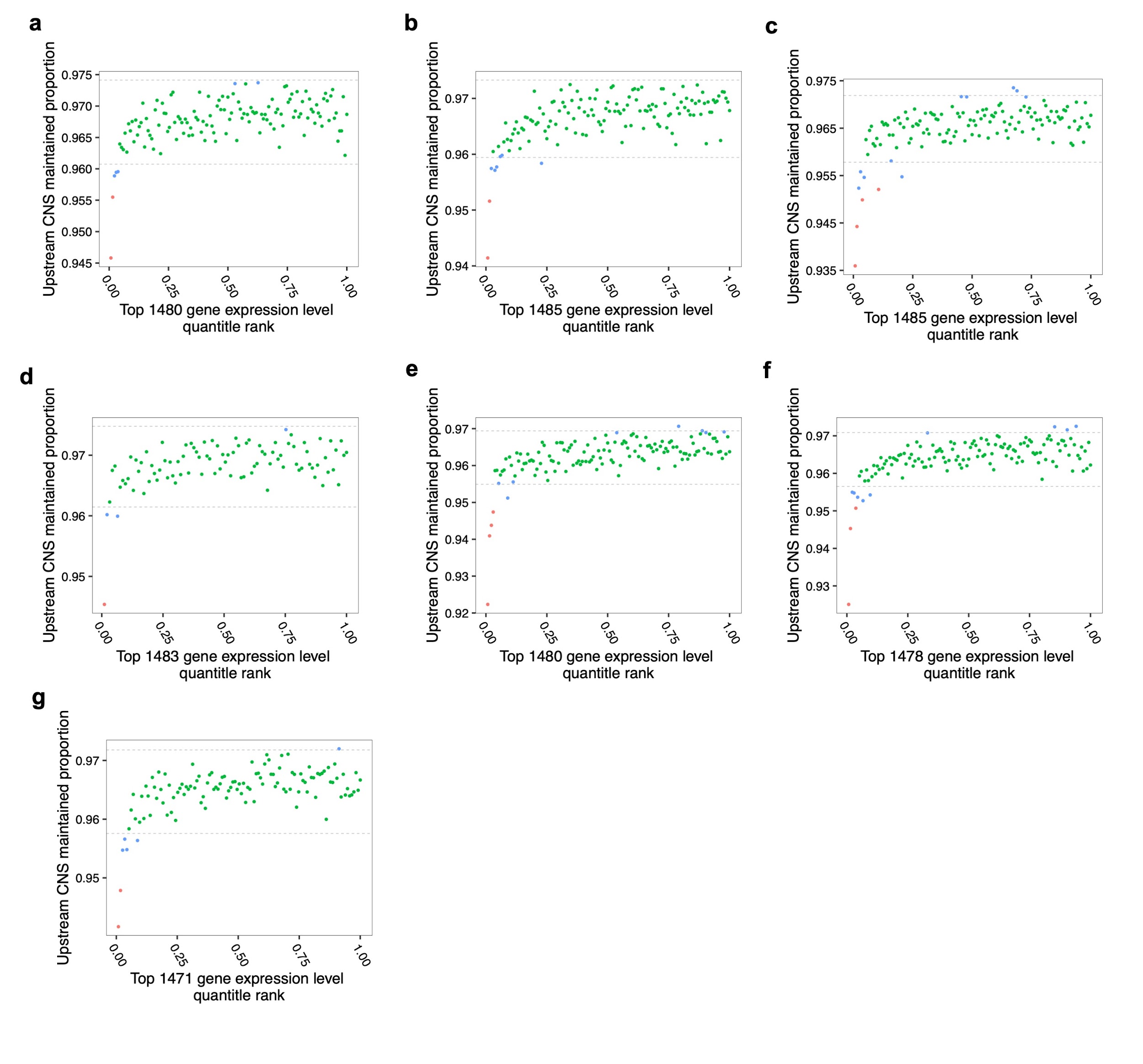

Sfig. 11 Comparing the proportion of maintained CNS in upstream 2Kb region of each accession and each gene of maize inbreeding population with the gene expression level rank of top 1500 high expressed genes. The gene expression level was quantified using B73 v3 reference and uplifted to v4 reference, and not all the genes could be uplifted to maize v4 reference annotation. **a,** Roots of germinating seedlings. **b**, Shoots of germinating seedling. **c**, Adult leaves collected at night. **d**, Adult leaves collected during the day. **e**, Tip of leaf three. **f**, Base of leaf three. **g**, Kernels at 350-growing-degree days.

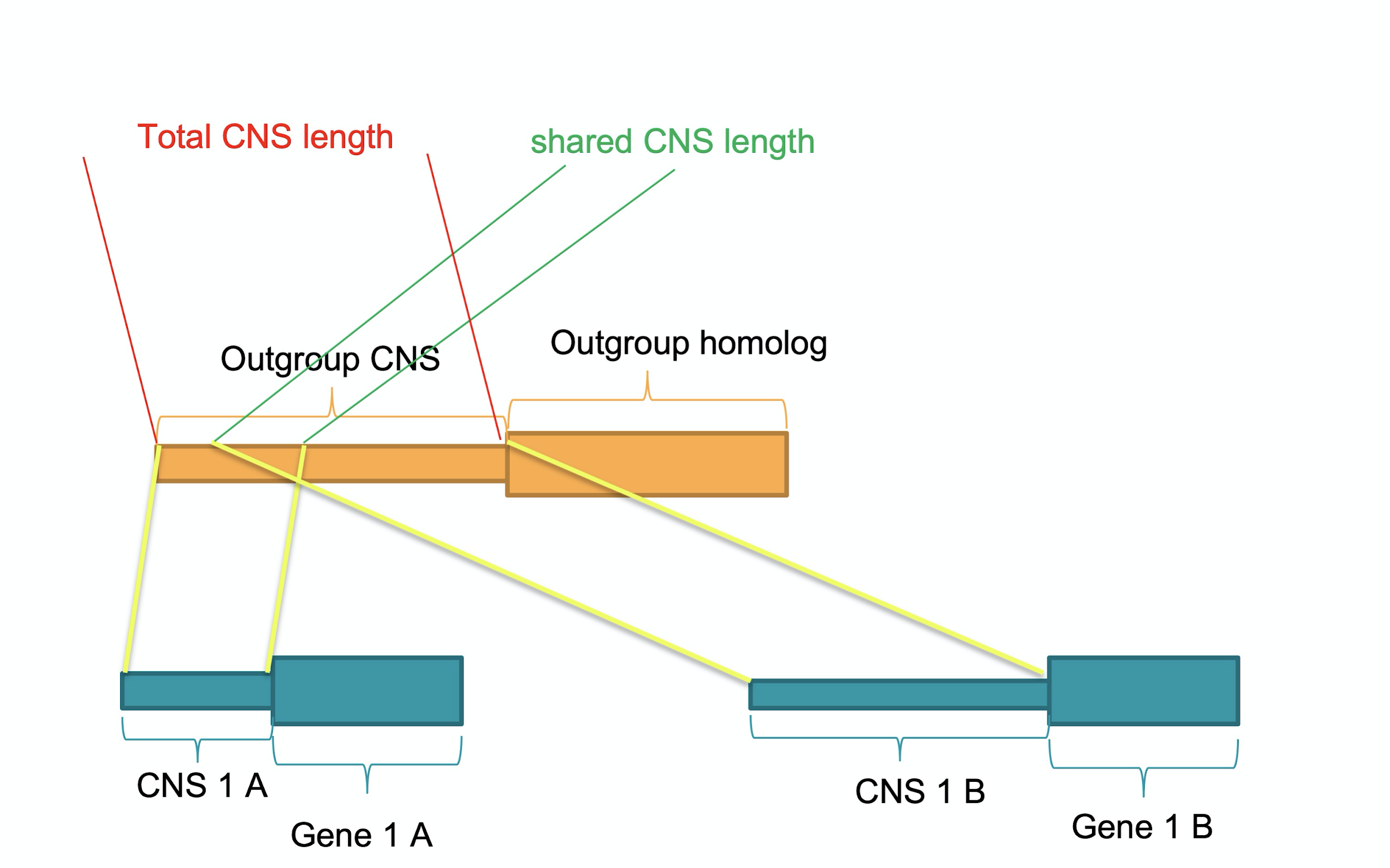

Sfig. 12 This cartoon shows how to calculate the CNS shared by two maize paralogous from different subgenomes using sorghum as the outgroup. We counted the number of bps shared between maize and sorghum in the 2Kb upstream of homolog, and counted how many bps appeared in each duplicated maize copy. For pan-And-CNS and core-And-CNS we released the condition that outgroup CNS should be within the 2Kb upstream of homolog.

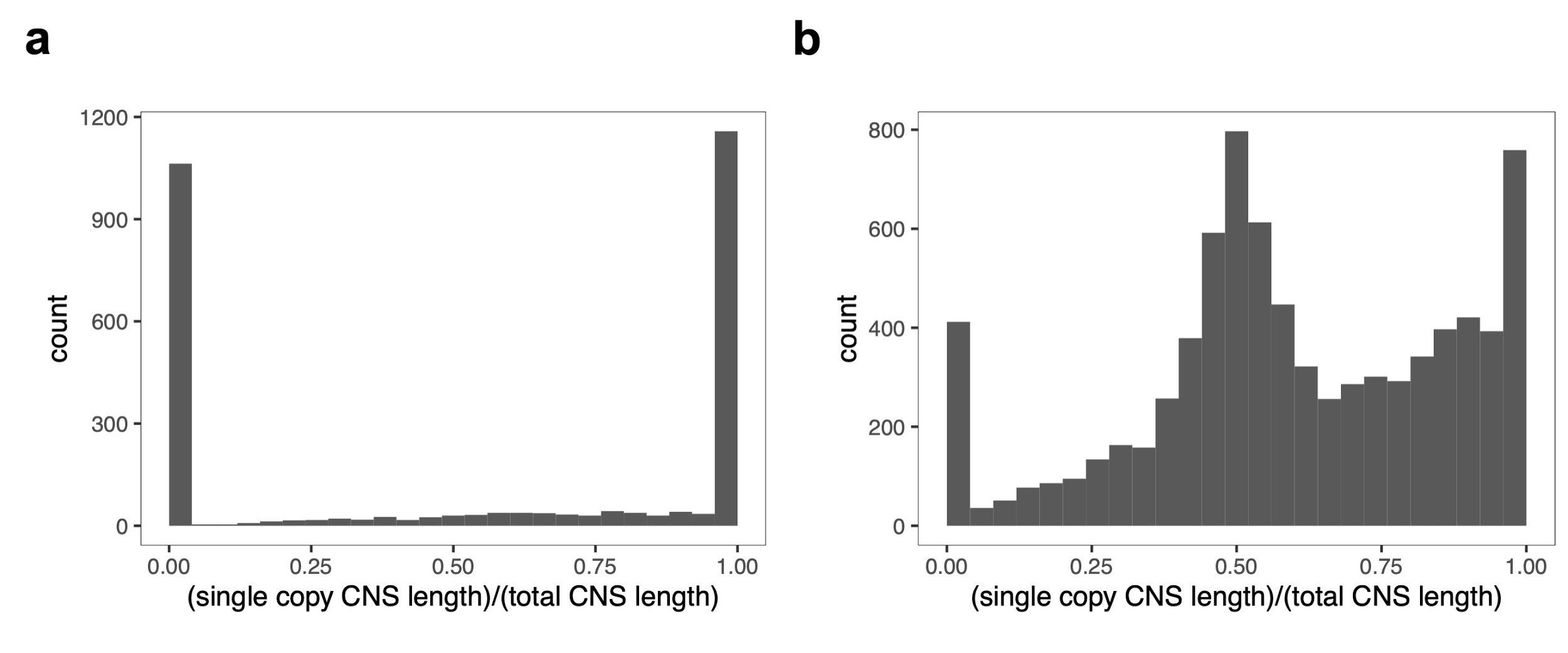

Sfig. 13 In the upstream 2Kb region of homologous gene pairs from different subgenomes of maize, we counted how many base-pair of core-and-CNS is present in each copy, how many base-pairs are shared and how many base-pair in total. And calculated the proportion of CNS present in each copy. The copy1 and copy2 were assigned arbitrarily. **a,** core-And-CNS result. **b,** pan-And-CNS.

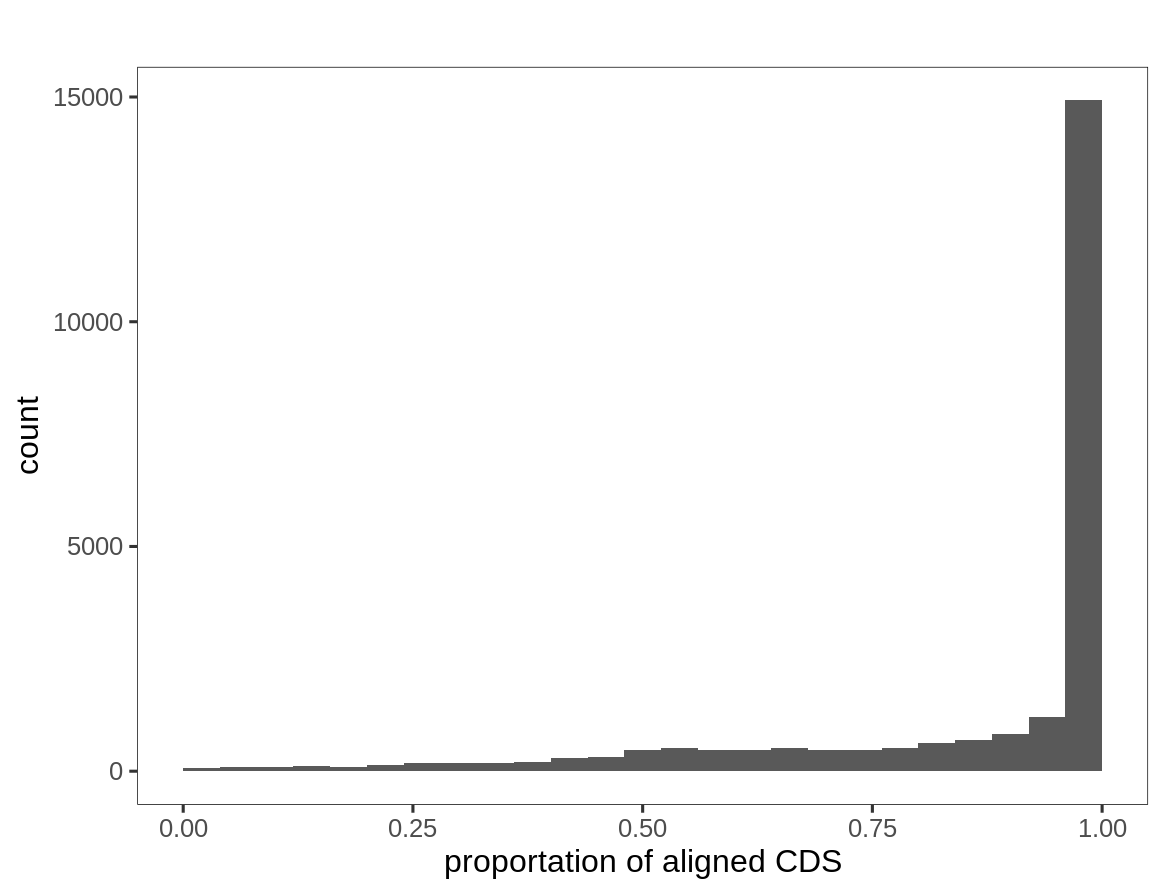

Sfig. 14 We counted how many base-pair of outgroup (sorghum) CDS is alignable in each copy of the mazie syntenic duplicated gene pair. And calculated the proportion of alignable in the CDS region.

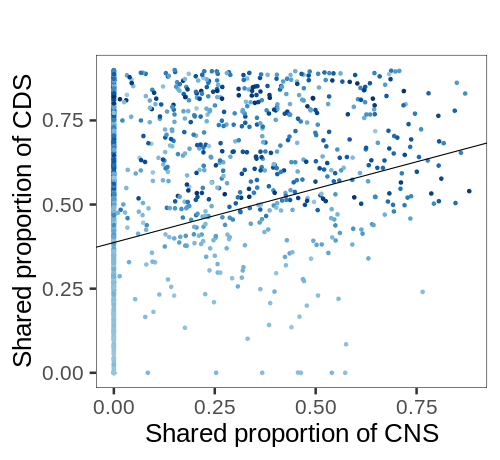

Sfig. 15 The portation of shared CNS in upstream 2Kb and CDS are positively correlated between the mazie syntenic duplicated gene pair. Only shared CNS <0.9 and shared CDS<0.9 are showing here.

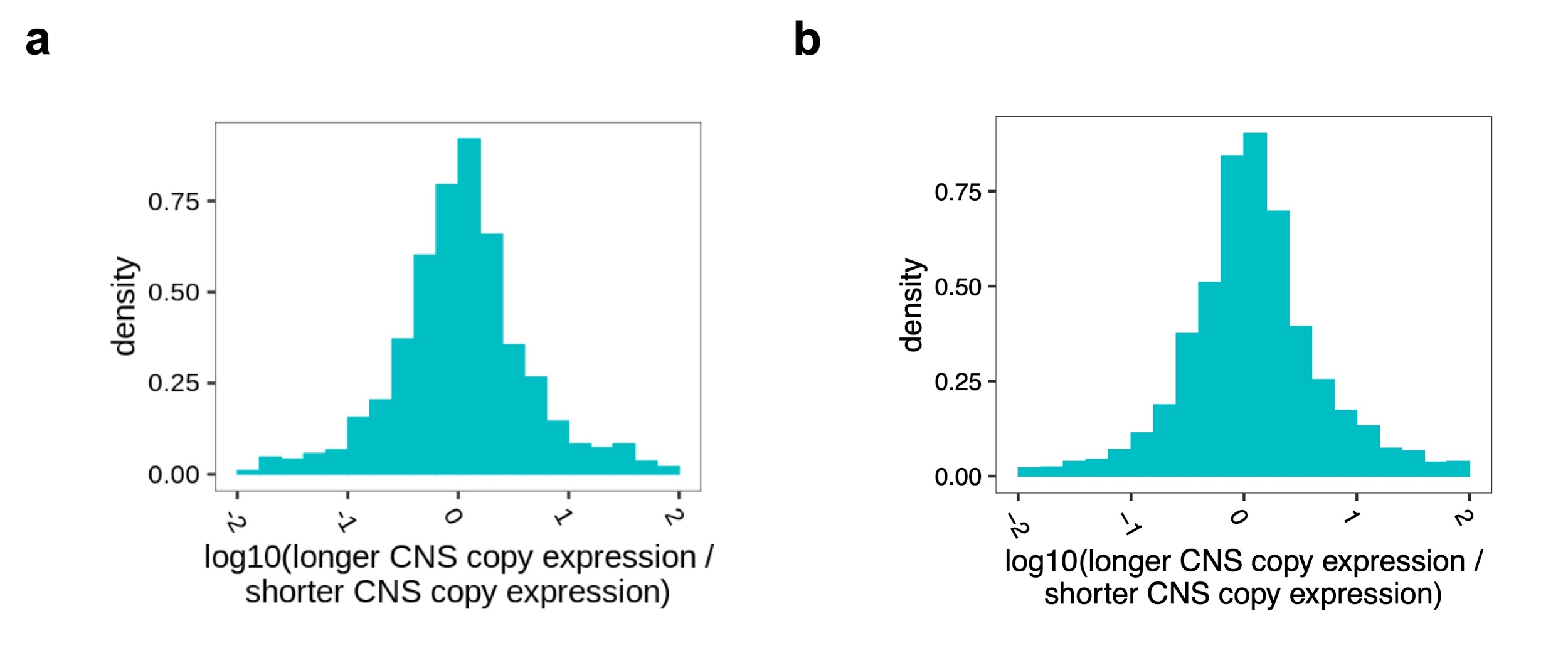

Sfig. 16 Between homologous gene pairs from different subgenomes of maize, the copy with longer core-And-CNS is associated with higher expression level. a, core-And-CNS result. b, pan-And-CNS result.

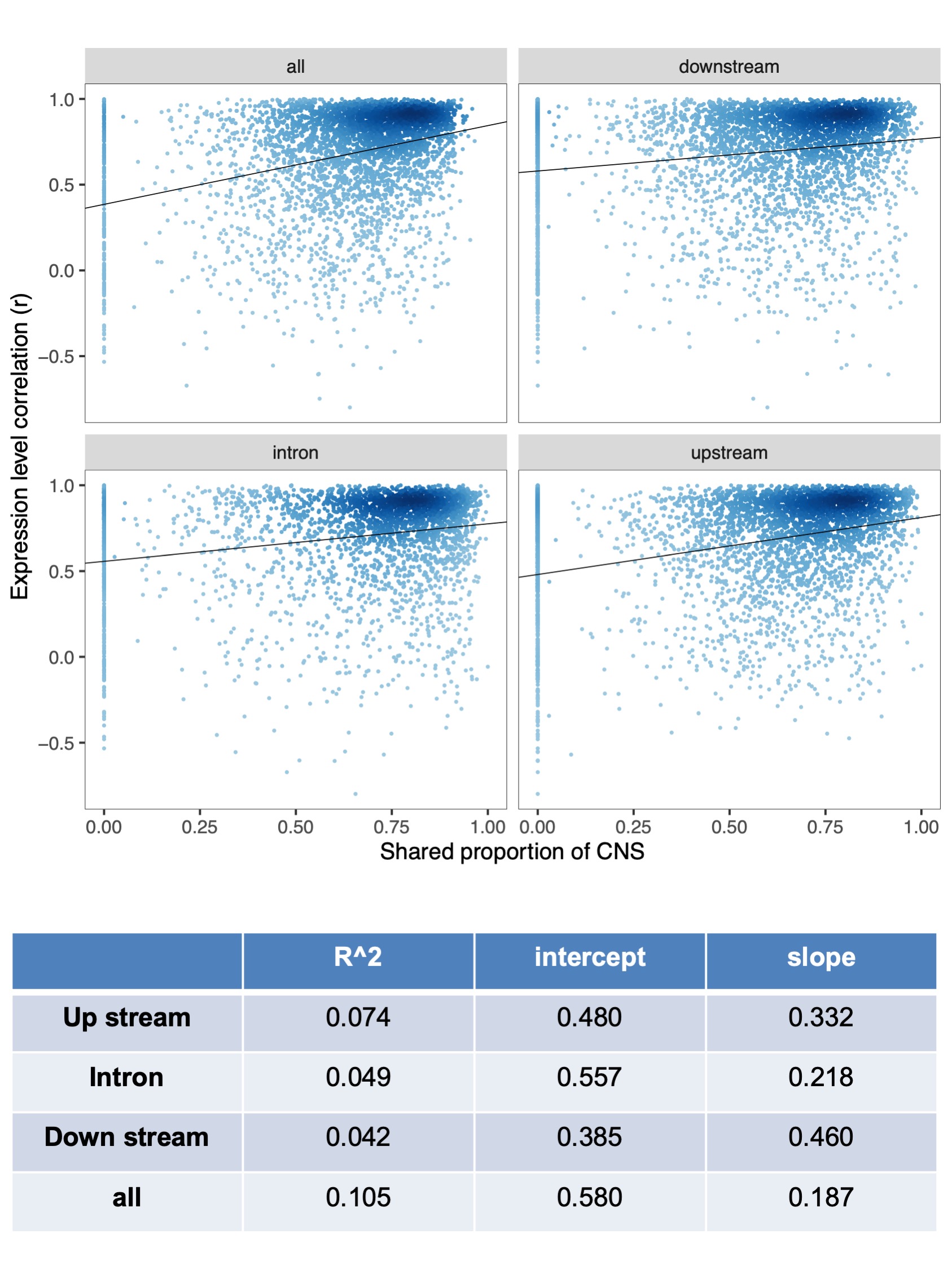

Sfig. 17 Comparing the CNS similarity with the expression level similarity of homologous gene pairs from different subgenomes of maize. **a,** X-axis is the proportion of shared CNS in upstream 2Kb range, downstream 2Kb range, intron region and overall the regions. And the y-axis is the expression level similarity of maize subgenome gene pairs measured using Persion correlation (r). **b,** the regression result between CNS similarity with the expression level similarity. The proportion of shared CNS was square transformed. CNS detected between maize and sorghum was used for this plot.

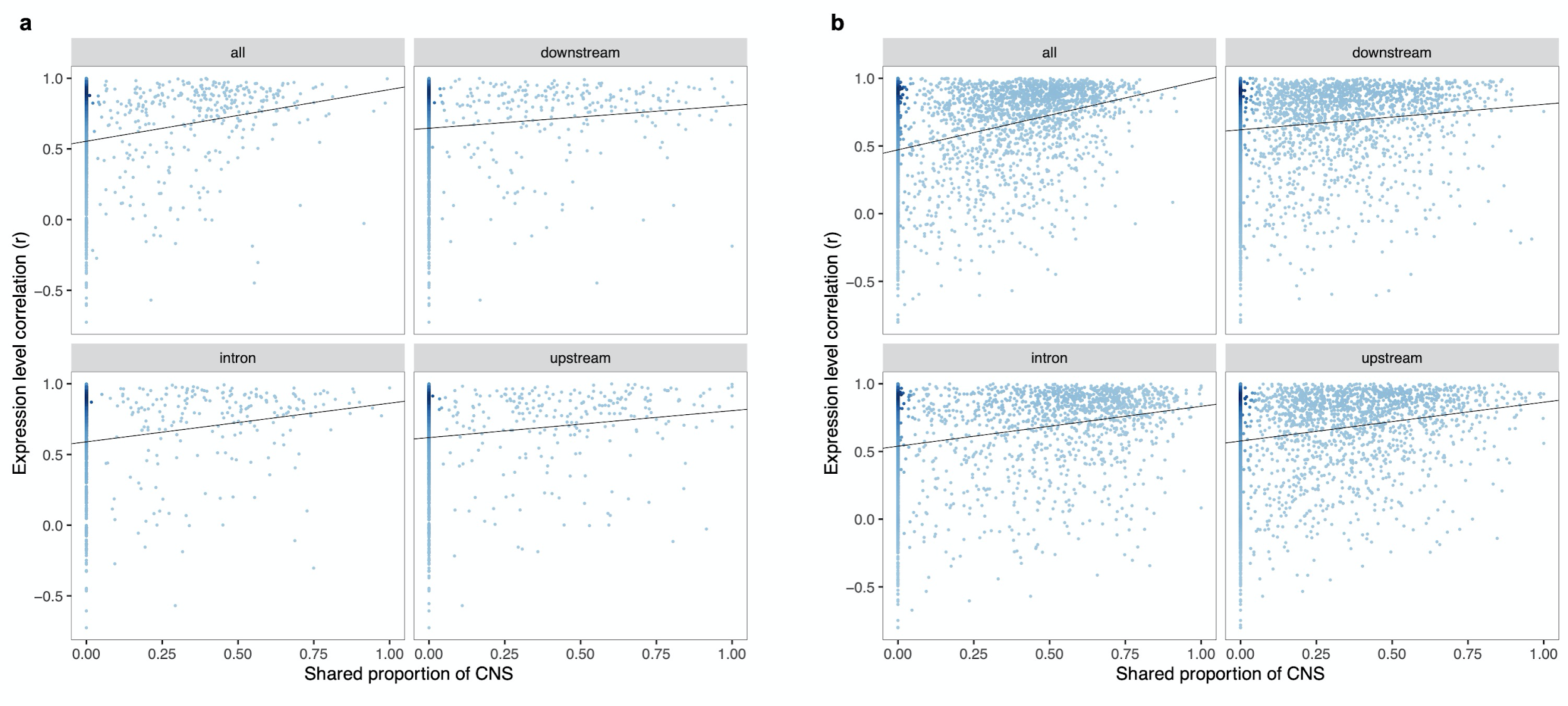

Sfig. 18 Comparing the CNS similarity with the expression level similarity of homologous gene pairs from different subgenomes of maize. **A,** plot using core-And-CNS. **b**, plot using pan-And-CNS.

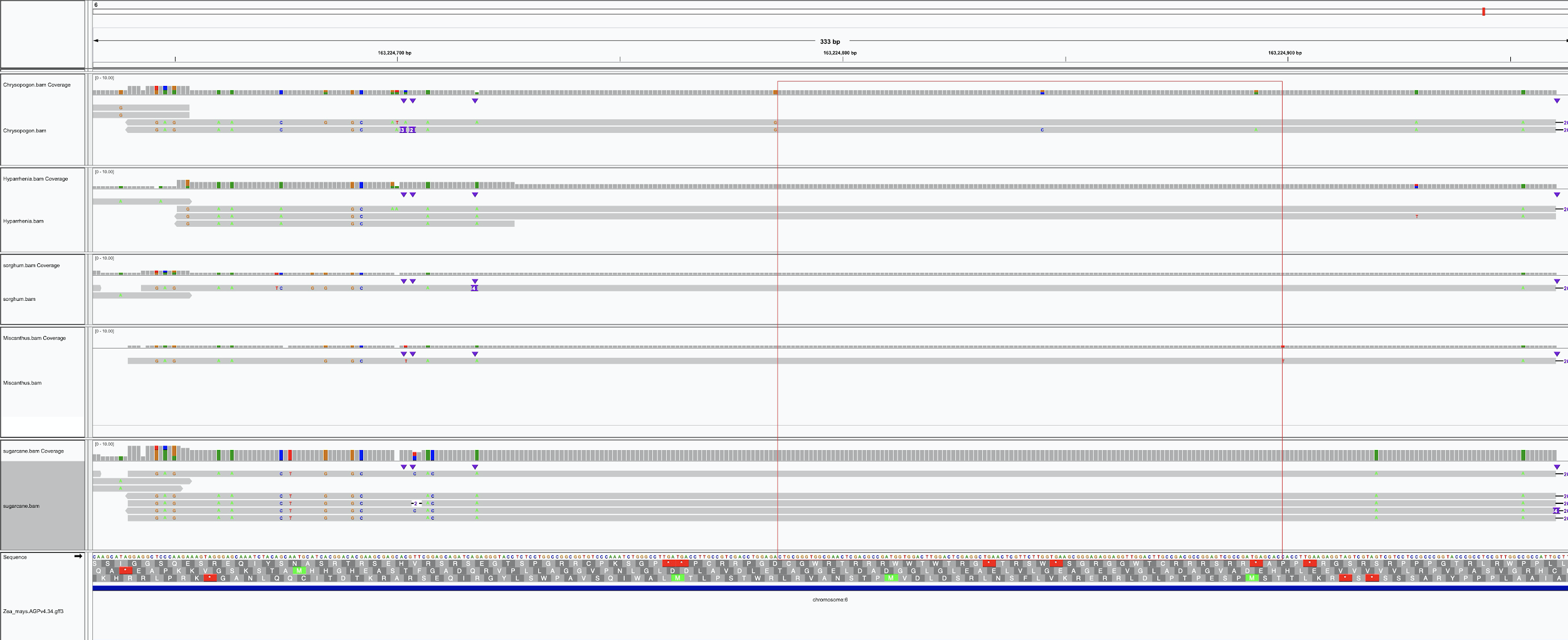

Sfig. 19 The sequence alignment in the region with the longest absolute CNS detected. The longest CNS is located in the intergenic region.

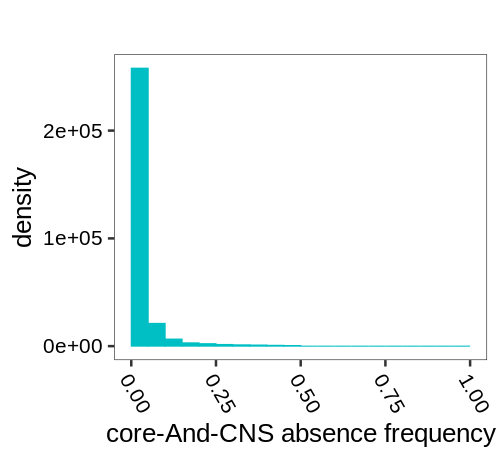

Sfig. 20 The absence frequency of core-And-CNS.

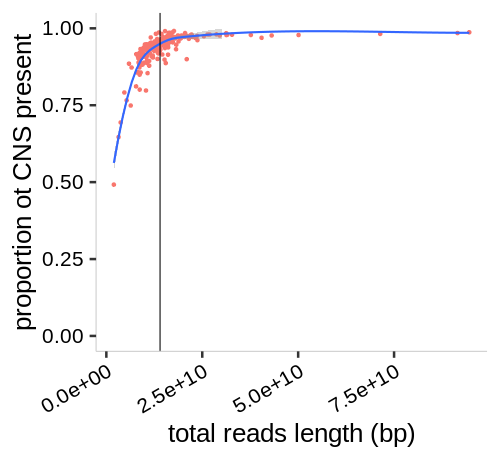

Sfig. 21 The number of CNS encoded as present is related to the sequencing scale in the maize population using whole genome sequence data. The x-axis is the total number of sequencing reads base-pairs. And the y-axis is the number of CNS encoded as present. Each dot represents a maize accession. And for each CNS in each accession, if the number of present base-pair is longer than 40% is CNS length in the maize reference genome, it is defined as presence, otherwise, absence.
